# Structure of Photosystem I with SOD reveals coupling between energy conversion and oxidative stress protection

**DOI:** 10.1101/2025.07.05.663305

**Authors:** Yue Feng, Kun Huang, Xiaobo Li, Alexey Amunts

## Abstract

In photosynthetic organisms, Photosystem I (PSI) not only converts energy into its chemical form but is also a major source of superoxide (O_2_^−^), which can damage cellular components. Its detoxification by superoxide dismutase (SOD) is essential for redox homeostasis. We report the structure of the PSI-SOD supercomplex from *Chromera velia* (*C. velia*), revealing a direct coupling between energy conversion and oxidative stress protection. The heterodimeric SOD comprises a catalytic SOD2 and a non-catalytic SOD1 anchoring it to the conserved stromal PSI surface near the ferredoxin-binding site with four contact areas. Structural comparison with plants and algae shows that SOD could occupy a similar position via interactions with PsaD. *C. velia* PSI displays several distinct features, including a split subunit PsaA linked by PsaV, a unique iso-fucoxanthin bound to PsaB, and an extended PsaF coordinating an extra chlorophyll in the membrane. The antenna includes five fucoxanthin-chlorophyll family proteins (FCPs) stabilized by adaptations unique to each antenna protein. These findings reveal the structural basis for the integration of a photoprotective mechanism with energy metabolism in photosynthetic membranes.

## Main

Photosynthesis enables cyanobacteria, algae, and green plants to capture solar energy for metabolism and carbon fixation. During electron transport, molecular oxygen can be inadvertently reduced, forming O₂⁻ ^1,2^. This reactive species damages macromolecules and lipids, contributing to chloroplast membrane rupture and cell death^3–6^. Due to its strong reducing power, PSI is a major generator of O₂⁻ on its stromal side^7,8^, from where it can diffuse and trigger oxidative cascades^9^. To mitigate oxidative stress, photosynthetic organisms rely on SOD that converts O₂⁻ into H₂O₂, subsequently removed by peroxidases^8^. SOD isoforms are found in the cytoplasm, chloroplasts, and mitochondria. In plants, over half of Fe-SOD proteins are attached to the stroma where PSI is localized^10,11^, and they play a role in protecting chloroplast nucleoids and metabolic pathways^12^. Thus, while PSI has been optimized as a highly efficient photoelectric machine^13,14^, the evolution of photosynthetic complexes has also prioritized photoprotection^15,16^. Previous studies estimated the local concentration of SOD in the vicinity of PSI is ∼1 mM^17^, and it is believed that PSI may physically recruit SOD for localized defense suggesting a mechanism for directly neutralizing PSI-produced O₂⁻ analogous to respiratory complexes^18^. The co-migration of SOD with PSI has been demonstrated by 2D gel electrophoresis and mass spectrometry^19^.

To capture the interactions of PSI with SOD, we used a chromerid alga *C. velia.* In these species, it was previously reported that SOD could be stably associated with PSI^19^. We purified the PSI-SOD-FCPI supercomplex (Extended Data Fig. 1a) and confirmed the presence of SOD subunits by SDS-PAGE and mass spectrometry (Extended Data Fig. 1b). High-performance liquid chromatography (HPLC) analysis showed that the supercomplex contains four major pigment species: chlorophyll *a* (Chl *a*), iso-fucoxanthin (iFx), violaxanthin (Vio), and β-carotene (Bcr) (Extended Data Fig. 1c). The material was then subjected to single-particle cryo-electron microscopy (cryo-EM) analysis, and the data were processed with cryoSPARC^20^. Upon *ab initio* reconstruction, 225,926 PSI particles were selected. We then performed a second step of classification based on the stromal region, retaining 88,960 PSI-SOD particles that were subsequently refined to 2.66 Å resolution. To further improve fragmented density in the peripheral regions, we iteratively refined the particles, selected 56,990 with additional density, and performed masked focused refinement on two antenna parts that yielded 2.77 Å and 3.03 Å reconstructions of the FCP proteins extending from core (Extended Data Fig. 2a, b; Supplementary Table 1). The complete model for PSI-SOD-FCPI was initially built using CryoFold^21^ and finalized manually with *Coot* 0.8.9^22^ and PHENIX^23^ (Extended Data Figs. 3 and 4; Supplementary Table 1).

The supercomplex model accommodates two SOD subunits and five peripheral light-harvesting FCP subunits (a, b, c, d, e) per PSI core (Fig. 1a, b). The PSI core consists of 12 subunits: PsaA1, PsaA2, PsaB, PsaC, PsaD, PsaE, PsaF, PsaI, PsaL, PsaM, PsaR, and the newly identified subunit PsaV (NCBI: taxid1169474) (Fig. 1a, b; Extended Data Fig. 3). Notably, PsaV is a lumenal subunit that links both PsaA segments and contributes structurally to PSI integrity. On the stromal side, the association of SOD1 and SOD2 with PSI is mediated by the extended PsaD and PsaM subunits (Fig. 2a; Extended Data Fig. 5). The cryo-EM map also revealed 12 ligands and 123 pigments, including 104 Chl *a*, 11 Bcr, seven violaxanthins and one unique iso-fucoxanthin designated as chromeraxanthin (Fig. 1c; Extended Data Fig. 1c; Extended Data Fig. 4; Supplementary Table 2).

**Fig. 1.**
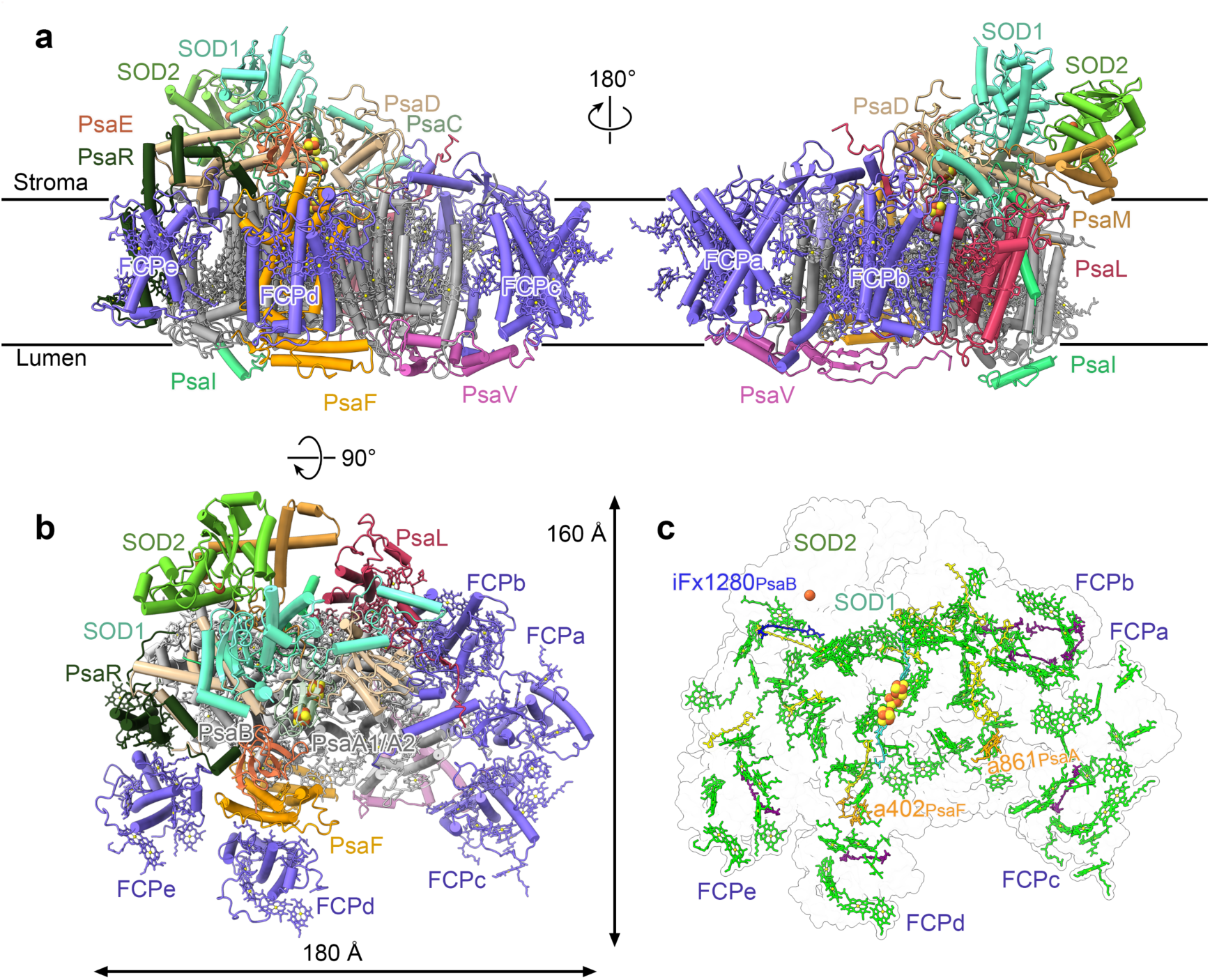
Overall structure of the PSI-SOD-FCPI supercomplex. **a**, Model of the PSI-SOD-FCPI supercomplex colored by individual subunits. **b**, Stromal view of the supercomplex showing the bound SOD1-SOD2 and five FCP light-harvesting proteins. **c**, Model of cofactors. Chl *a* green, Bcr yellow, iFx blue, Violaxanthin purple, the chromeraxanthin and unique Chl *a*402_PsaF_ and Chl *a*861_PsaA_ are labeled.

**Fig. 2.**
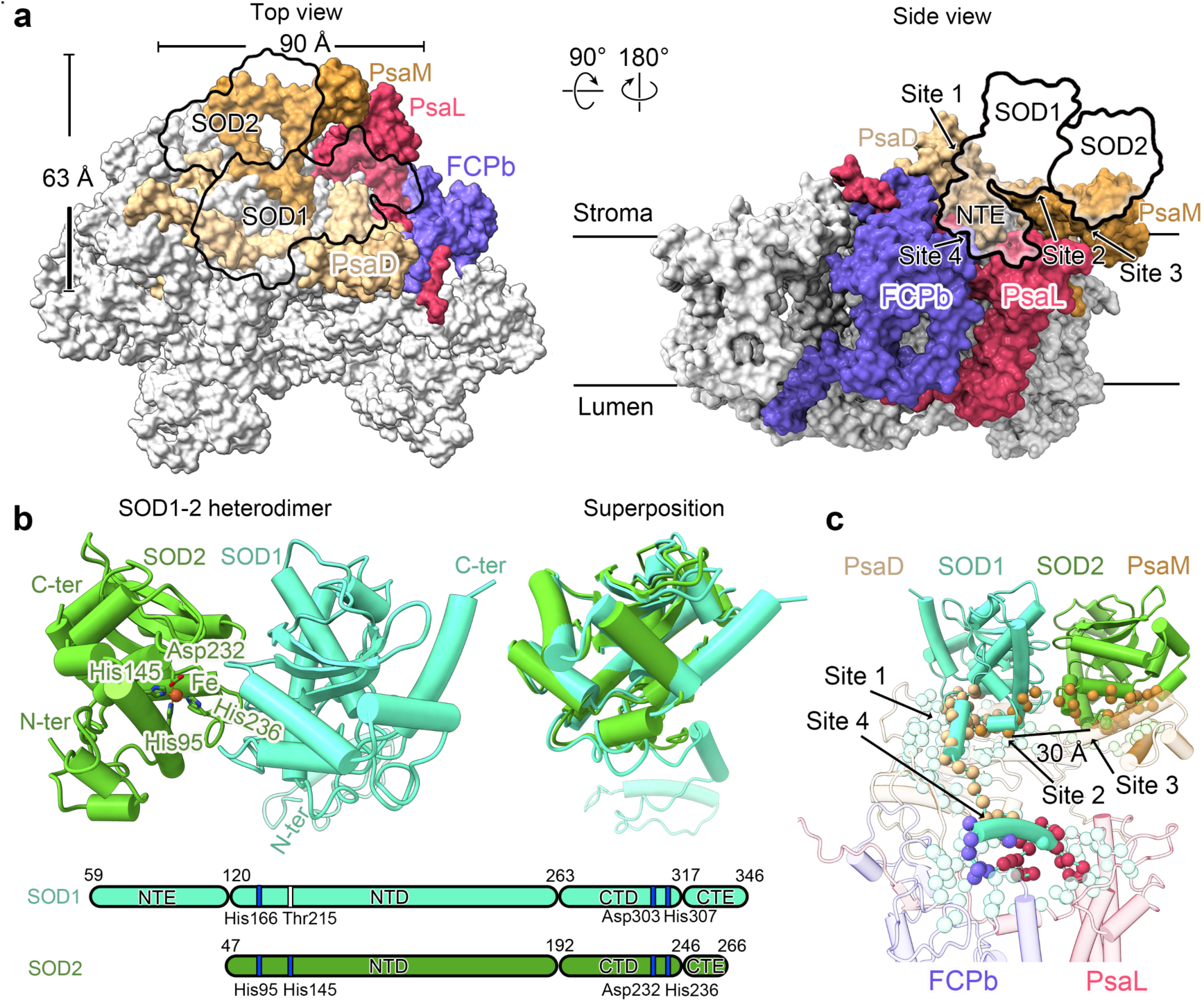
Binding of SOD1-SOD2. **a**, SOD1 and SOD2 are outlined, PSI-FCPI protein subunits that bind SOD in four contact sites are colored. **b**, Left, structure of the SOD1-SOD2 heterodimer with cofactor Fe and its coordinating residues. Right, superposition of SOD1 and SOD2 shows the N-terminal extension. **c,** Interacting residues between SOD1-SOD2 and PSI-FCPI are shown as spheres. SOD residues are colored consistently with the subunits they interact with. The four contact sites are indicated.

In our PSI-SOD-FCPI supercomplex, the SOD complex is organized as a heterodimer, with SOD2 is a 24-kDa subunit that binds catalytic iron (Fe), coordinated by His95, His145, Asp232, and His236. SOD1 is a 32-kDa subunit with an N-terminal extension (NTE) of 61 residues that stabilizes interactions with PSI (Fig. 2a,b) and extends toward the ferredoxin binding site (Extended Data Fig. 6). Together, the SOD1-SOD2 module forms a stromal complex on PSI, consistent with its function as a first-line defense against oxidative stress^24^.

The connection between SOD and PSI involves four contact sites on the stromal side, involving subunits PsaD, PsaL, PsaM, and FCPb comprising a total buried surface area of ∼5180 Å^2^ (Fig. 2a, c; Supplementary Table 3). Site 1 involves the NTD of SOD1 interacting with the conserved region of PsaD (residues 36 to 148). This highly conserved interface forms a tight association and is likely preserved in PSI across various organisms. Site 2 is formed by the NTD of SOD1 contacting the C-terminal end of PsaM. This interaction is part of a pseudosymmetric arrangement with Site 3 and is stabilized by several specific residue-level contacts. Site 3 mirrors Site 2, involving the NTD of SOD2 and forming similar contacts with a second region of the PsaM C-terminus. The two sites are separated by ∼30 Å, reflecting the heterodimeric configuration of the SOD complex. Site 4 is formed by the NTE of SOD1 and undergoes a sharp 90° bend that runs along the stromal surface of PSI. We were able to model 61 residues within this extension, which adopts a conformation composed of two α-helices (Fig. 2b). The NTE slides along and interacts with stromal-facing regions of three PSI components: PsaD (residues 36-148), PsaL (residues 52-103), and FCPb (residues 170-194), providing additional stabilization of SOD at the periphery of the supercomplex (Fig. 2c).

The binding of the SOD1-2 protein heterodimer is compatible with the presence of the soluble electron acceptor ferredoxin on PSI^25^, suggesting that no conformational changes are required for dealing with the oxidative damage (Extended Data Fig. 6). In the event of P700 over-reduction and electron leakage in PSI, the bound SOD can directly disproportionate superoxide radicals into hydrogen peroxide and oxygen, in line with the pseudo-cyclic electron transport pathway known as the water-water cycle (Mehler reaction)^1^. Since the binding site for ferredoxin is conserved^25,26^, the association of SOD may also be conserved, but more stable in *C. velia* due to protein extensions.

Structural comparison with PSI from plants^27,28^ and green algae^29^ shows that SOD could occupy a similar position in the supercomplex as observed in our structure, with no apparent steric clashes through PsaD interactions (Extended Data Fig. 7). Thus, PSI-SOD supercomplex may not be a unique feature of *C. velia*. Indeed, plant homologs of FeSODs, namely FSD2 and FSD3 also form a heterocomplex and its disruption leads to the accumulation of reactive oxygen in *Arabidopsis*^12^. Our results further suggest a structural basis for why FSD2 and FSD3 are not functionally interchangeable^30^. In the PSI-SOD structure, the binding of SOD1 and SOD2 to the PsaD side is pseudosymmetric with SOD1 acting as an anchor for the heterodimer, highlighting structurally distinct roles for the two subunits.

From broader perspective, protein crowding within the membranes of bioenergetic organelles is an emerging theme elucidated with recent structural studies^31^, PSI is often found associated in higher oligomeric states^29,32^. In mitochondria, immunoblotting of *Caenorhabditis elegans* respiratory supercomplexes separated by BN-PAGE showed that SOD-2 is associated with the respirasome, and in Gram-positive bacteria, the catalytically active SodC is an integral part of a respiratory supercomplex^18^. Thus, the presence of PSI-SOD supercomplex may facilitate a more effective and compartmentalized defense against oxidative damage by enabling electron abstraction directly from ROS generators during photosynthesis.

The PSI core of *C. velia* shares general similarity with that of *Symbiodinium sp.*^33,34^, including Qy absorption and 77 K emission peaks at 679 nm and 708 nm, respectively (Extended Data Fig. 8). However, we identified several distinct and functionally important structural features that modulate the energy transfer. One of the most notable differences is the splitting of the central subunit PsaA into two separate polypeptides, resulting in the N-terminal PsaA1 (20 kDa) and the C-terminal PsaA2 (42 kDa). On the stromal side, a 20-residue loop is absent, disrupting the continuity of the protein backbone (Fig. 3a). This structural gap is compensated by the newly identified subunit PsaV, which binds to both PsaA1 and PsaA2 and restores connectivity (Fig. 3a). A new chlorophyll molecule, Chl *a*861, is inserted between the split domains, to mediate the energy transfer pathway from PsaA1 to PsaA2 via Chl *a*820 and Chl *a*830 with Mg-Mg distances of 24.6 Å and 16.6 Å, respectively (Fig. 3a). Since chlorophyll biosynthesis and membrane insertion are synchronized with the biogenesis of photosynthetic proteins ^35^, the integration of Chl *a*861into independently synthesized PsaA1 and PsaA2 by the chlororibosome ^36^ is likely to be coordinated.

**Fig. 3.**
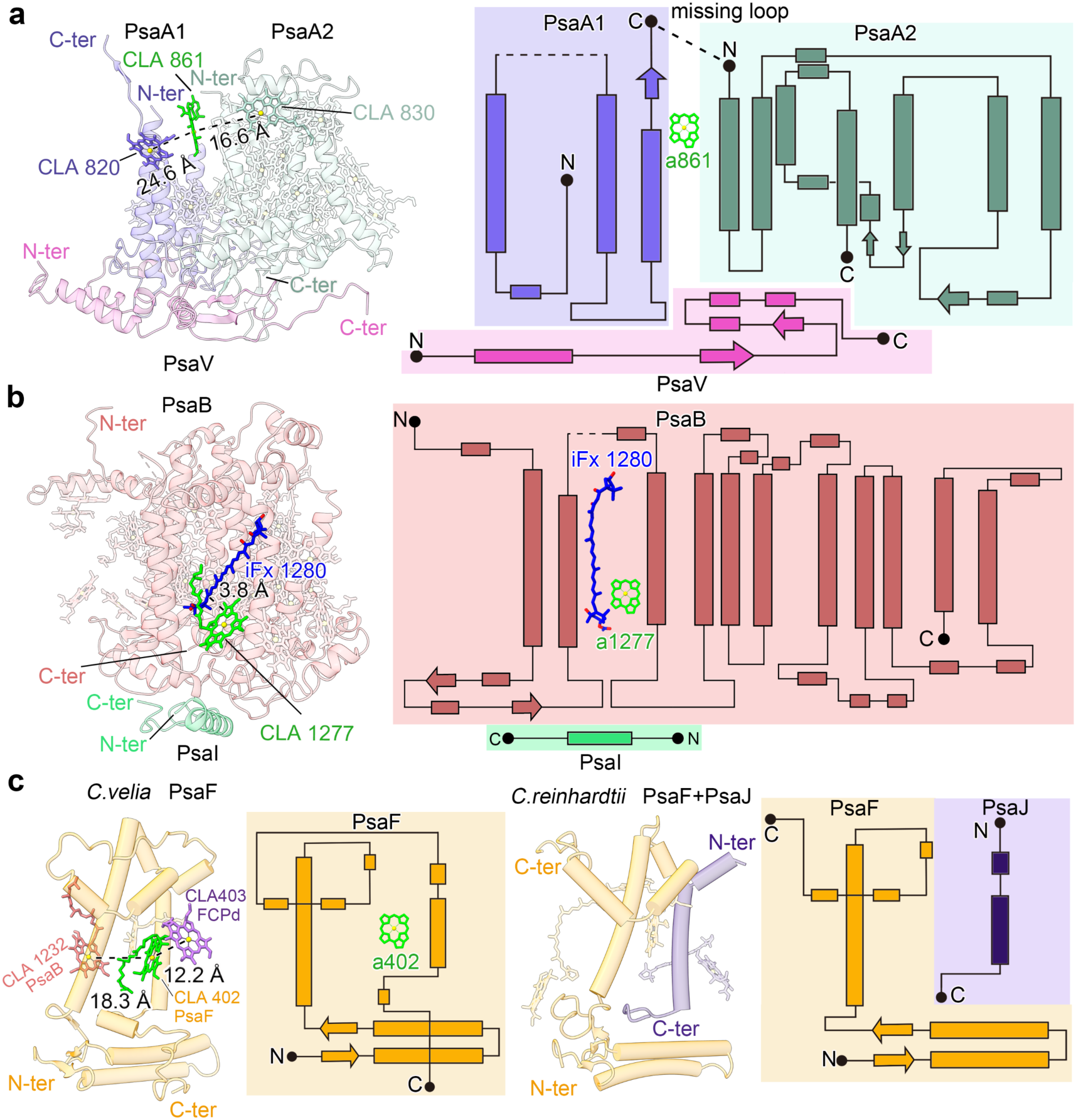
Distinct features of *C. velia* PSI. **a**, Split subunit PsaA with a new Chl *a*861 between the two segments. The topology diagram shows the organization of the domains with PsaV linking PsaA1 and PsaA2. **b**, Structure and topology diagram of subunit PsaB stabilized with the luminal PsaI and coordinated chromeraxanthin with associated Chl *a* 1277. **c**, The extended PsaF (left) replaces the canonical PsaJ (right) and coordinates an extra Chl *a*402 that mediates an additional energy transfer pathway by bridging FCPd to PsaF and PsaB.

In the PsaB subunit, a lumenal extension is sealed by the terminal α-helix of PsaI, where our cryo-EM map revealed an elongated non-protein density absent in other known PSI structures (Extended Data Fig. 9a,b). The density, approximately 33 Å in length is associated with Chl *a*1277 (Extended Data Fig. 9a). Guided by previously published PSI-FCPI structures^37–40^, we modeled a fucoxanthin molecule in the density (Extended Data Fig. 9b). However, the density in its second ring differs, corresponding to a cyclopentane instead of the canonical epoxy group found in typical fucoxanthin, a configuration not matching any previously known iFx isoforms (Extended Data Fig. 9b). This distinct feature identifies it as a novel isoform, designated chromeraxanthin. Positioned between the second and third transmembrane helices of PsaB (Fig. 3b), its acetyl group extends into the hydrophobic gap between lipid molecules, while its conjugated polyene chain is closely surrounded by the phytol chain of Chl *a* 1277 (Extended Data Fig. 9a). The acetyl group end contains an adjacent allenic bond that is typical of the fucoxanthin family. Biosynthesis of this moiety from the violaxanthin precursor has been partially elucidated, involving violaxanthin de-epoxidase-like enzymes and a putative acetyltransferase^41,42^. Chromeraxanthin is abundant in the peripheral antenna pool^43^, and shows effective green-blue light (450-550 nm) absorption capabilities (Extended Data Fig. 1c). The chromeraxanthin is coupled to Chl *a*1277 within 3.8 Å, which is typical to fucoxanthin-Chl *a* interactions in diatom^44^. This enables π–π stacking interactions, facilitating efficient Förster resonance energy transfer (FRET), suggesting an alternative excitation energy transfer. Recent report on the influence of keto and hydroxy functional groups on energy transfer further suggests a possible fine tuning of energy funneling^45^.

Finally, the canonical transmembrane subunit PsaJ is absent in *C. velia*, and its role appears to be assumed by an 81-residue C-terminal extension of PsaF, the subunit that is responsible for plastocyanin binding. The extended PsaF traverses the membrane (from the stroma to the lumen) where it coordinates an extra Chl *a*402 and reaches the plastocyanin binding site (Fig. 3c). This chlorophyll mediates an additional energy transfer pathway by bridging FCPd to PsaF and PsaB via close pigment-pigment distances (Chl *a*403, 12.2 Å; Chl *a*1232, 18.3 Å). Notably, a fused *psaF-psaJ* sequence has previously been identified in a marine viral genome^46,47^. Together, these structural innovations introduce three previously unreported cofactors: two additional chlorophylls and the unique chromeraxanthin, arising from specific protein rearrangements that support alternative excitation energy transfer pathways within PSI-FCP.

The PSI-FCP supercomplex features a peripheral antenna composed of five distinct fucoxanthin-chlorophyll proteins (FCPa-e), each adopting unique structural features resolved in our 2.77-3.03 Å local resolution map. This reflects an adaptation to the remodeled PSI core surface (Fig. 4; Extended Data Fig. 10). FCP structures are generally conserved in the core, having three transmembrane helices, but differ from one another in protein length (160-247 residues) and pigment content (7-12 chlorophylls, 0-4 fucoxanthins). FCPa, FCPc, FCPd, and FCPe belong to the Lhcr family with FCPa being extended with a stromal α-helix that docks it to the core at the PsaA pole; FCPb is phylogenetically independent and belongs to a distinct monophyletic clade CgLhcr9^40,48^ (Extended Data Fig. 11). Each FCP has been structurally adapted to interact with a different PSI core subunit: FCPa-PsaA1, FCPb-PsaA2, FCPc-PsaA1/PsaV, FCPd-PsaF, FCPe-PsaR (Extended Data Fig. 10), indicating co-evolution to maintain efficient energy transfer despite PSI core’s architectural divergence. In addition, FCPa-b-c contact each other forming a hetero-trimer (Fig. 4; Extended Data Fig. 10). Computational analysis of PSI-SOD-FCPI based on FRET model^49^ shows the energy transfer pathways from each FCP to the PSI core subunit (Extended Data Fig. 12, Supplementary Table 4).

**Fig. 4.**
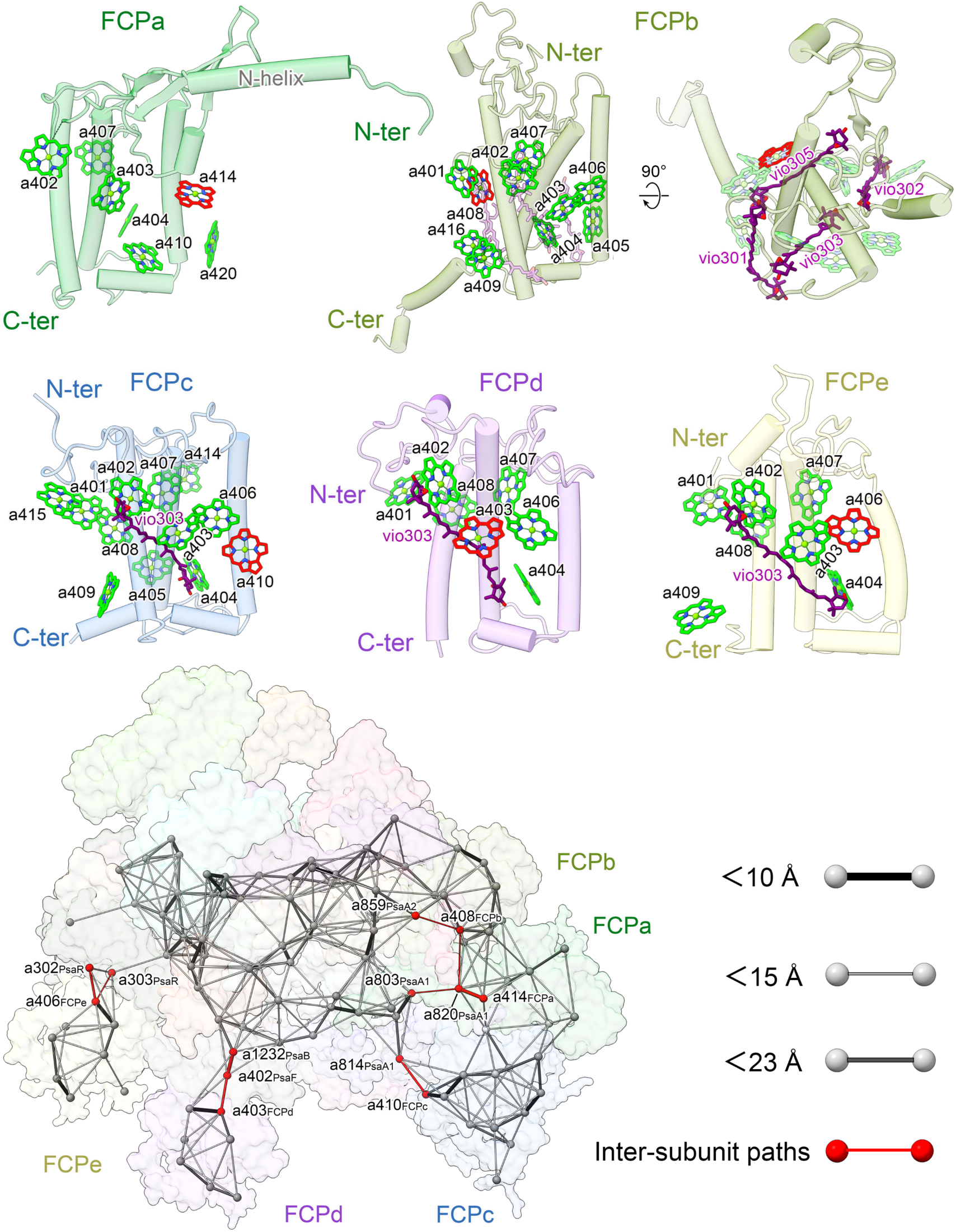
FCP light-harvesting proteins and energy transfer pathways. Individual FCPs are shown with the cofactors (phytol chains removed for clarity). Each FCP protein contains a conserved core but displays unique structural adaptations that enable specific interactions with different subunits of the PSI core. These adaptations include variable N- or C-terminal elements and loop regions that mediate contact with distinct PSI components. As a result, individual FCPs adopt diverse orientations around the PSI core and form specific functionally coupled chlorophylls for intersubunit energy pathways (Top panels, red chlorophylls). Bottom panel, energy transfer pathways with chlorophylls indicated by Mg atoms.

FCPa has the highest degree of structural specialization, featuring an N-terminal region extended by 110-residue domain forming an α-helix and three β-pleated sheets that engage the C-terminus of PsaA1 (Fig. 4; Extended Data Fig. 10). Notably, FCPa binds only seven Chls *a* and lacks carotenoids entirely (Supplementary Table 5). Functionally, its Chl *a*414 forms a coupled pair with Chl *a*820 of PsaA1, enabling energy transfer toward Chl *a*803 from a structurally distinct site (Fig. 4; Extended Data Figs. 10a, 12).

FCPb binds ten Chls *a* and four violaxanthins (Fig. 4; Supplementary Table 5). Its stromal extended N-terminal loop forms interactions with PsaD and PsaL and further stabilizes SOD1 (Extended Data Fig. 10). On the lumenal side, the C-terminal helix extends to PsaA2. We resolved four associated lipids, three of which are at the interface (Extended Data Fig. 10). Energy pathways are conducted through Chl *a*408 that is coupled to Chl *a*820 and Chl *a*859 of PsaA1 and PsaA2, respectively (Fig. 4, Extended Data Fig. 12).

FCPc interacts with the N-terminus of PsaV, that bridges the split PsaA1 and PsaA2. This association suggests that FCPc co-evolved with PsaV to maintain a continuous moiety in this newly formed configuration. Its Chl *a*410 funnels energy to Chl *a*814 of PsaA1 (Fig. 4, Extended Data Fig. 12). FCPd binds near the loop region of PsaF on the luminal side, an adaptation not seen in canonical PSI structures. The interactions include an extra Chl *a*402 from PsaF and Chl *a*406 from FCPd at their interface enabling a distinct energy transfer bridge to PsaB (Fig. 3c; Extended Data Figs. 10, 12). Finally, FCPe associates with PsaR, a minor PSI subunit, and transfers excitation via Chl *a*406 through Chl *a*302 and a303 of PsaR (Extended Data Figs. 10, 12).

Collectively, these observations reveal that each FCP has evolved a structurally distinct interface to complement the local topology and pigment architecture of its PSI docking site. The variability in orientation towards the core, coupled chlorophylls with PSI subunits, protein length, and surface contacts reflect selective pressures to maintain optimal light harvesting and energy transfer in the context of a remodeled PSI core scaffold. Thus, the architectural diversification of the light-harvesting protein family is driven by the need to adapt to specific PSI core surfaces for functional energy transfer.

In conclusion, the structure of the PSI-SOD-FCPI supercomplex reveals a direct spatial integration between energy conversion and oxidative stress protection. This association is mediated primarily via the NTD of SOD1 and the conserved region of PsaD, positioned near the ferredoxin-binding site, and is further stabilized by the NTE of SOD1 interacting with peripheral components. The stable placement of the SOD heterodimer adjacent to the electron transfer reaction suggests a conserved strategy to mitigate photodamage. Together with newly identified cofactors that modulate energy pathways and specialized FCPs, these findings underscore the evolutionary tuning of photosynthetic machinery to balance high-efficiency energy capture with localized photoprotection.

## Methods

### Purification of PSI-SOD-FCPI

*C. velia* CCMP2878 was obtained from the National Center for Marine Algae and Microbiota, USA. Cells were cultured at 25 °C under a photon flux density of 120 μmol photon m^-2^s^-1^ in the medium consisting of seawater, 8.82x10^-4^ M NaNO_3_, 3.62x10^-5^ M NaH_2_PO_4_H_2_O, 1.06x10^-4^ M Na_2_SiO_3_9H_2_O, 1.17x10^-5^ M FeCl_3_6H_2_O, 1.17x10^-5^ M Na_2_EDTA2H_2_O, 3.93x10^-8^ M CuSO_4_5H_2_O, 2.60x10^-8^ M Na_2_MoO_4_2H_2_O, 7.65x10^-8^ M ZnSO_4_7H_2_O, 4.20x10^-8^ M CoCl_2_6H_2_O, 9.10x10^-7^ M MnCl_2_4H_2_O, 2.96x10^-7^ M thiamine HCl, 2.05x10^-9^ M biotin, 3.69x10^-10^ M cyanocobalamin. All the following procedures were performed under green light and at 4 ℃ or on ice. Cells were harvested through centrifugation at 3,000*g* and resuspended in buffer containing 50 mM 2-morpholinoethanesulfonic acid (MES)-NaOH (pH 6.5), 5 mM MgCl_2_, 5 mM CaCl_2_. The pellet was resuspended and cells were disrupted by liquid nitrogen freezing grinding. The material was centrifuged at 1,000*g* for 10 min to remove unbroken cells. The supernatant was centrifuged at 150,000*g* for 30 min, and the pellet was resuspended in the buffer containing 50 mM MES-NaOH (pH 6.5), 10 mM NaCl, 5 mM CaCl_2_ to a concentration of 0.3 mg/ml chlorophyll and solubilized with 0.55 % (w/v) N-dodecyl-α-D-maltoside (α-DDM) (Anatrace, Maumee, OH, USA). Following 30-min incubation on ice, the material was centrifuged at 40,000*g* for 10 min and applied onto a continuous sucrose density gradient of 0.1-1.0 M (0.03% α-DDM) and centrifuged at 250,000*g* for 18 h in SW41 rotor (Beckman Coulter). The PSI-SOD-FCPI fraction was collected by syringe, sucrose was removed by dilution with the same buffer without sucrose, and the material was concentrated to 2.0 mg/ml chlorophyll.

### Characterization of the PSI-SOD-FCPI supercomplex

The subunit composition of the PSI-SOD-FCPI was analyzed by SDS-PAGE using a gel containing 14% polyacrylamide and 7.5M urea^50^. The gel was stained with Coomassie brilliant blue R-250 (Sigma-Aldrich, Germany). Protein identification was carried out by mass spectrometry. Briefly, Coomassie-stained bands were excited, subjected to in-gel digestion using sequencing-grade modified trypsin, and the resulting peptides were extracted and analyzed by mass spectrometry.

To investigate the gene families of each antenna subunit, the sequence of the five FCPIs resolved in our structure were compared with representative LHCs from typical red-lineage organisms, including *Chaetoceros gracilis* and *Thalassiosira pseudonana*^37,38^. Multiple sequence alignment was performed using CLUSTALW, and the phylogenetic tree was generated and visualized using iTOL v6.

The room-temperature absorption spectra and 77 K and fluorescence emission spectra were recorded using an ultraviolet-visible spectrophotometer (Shimadzu, Japan) and a fluorescence spectrophotometer (F-7000 Hitachi, Japan), respectively. Measurements were performed on the PSI-SOD-FCPI supercomplex at a concentration of 50 μg Chl/mL, wth fluorescence excitation at 436 nm.

Pigment composition was determined by HPLC as previously described^34^. The sample collected from the sucrose density gradient was mixed with 90% (v/v) acetone for 30 min at 4°C to extract the pigments. After centrifugation at 12,000*g* for 15 min, the supernatant was applied to a C8 column (150 × 4.6 mm, 5 μm particle size; GL Sciences, Japan), and the pigments were eluted at room temperature at a flow rate of 1 mL/min in linear gradient of buffer A (methanol: acetonitrile: 0.25 M pyridine = 50:25:25) transitioning to buffer B (methanol: acetonitrile: acetone = 20:60:20). Pigment identity was determined based on absorption spectra and retention times monitored at 445 nm. The results revealed the presence of violaxanthin, chromeraxanthin, Chl *a* and Bcr in the PSI-SOD-FCPI supercomplex.

### Cryo-EM data acquisition and processing

PSI-SOD-FCPI 2 mg/ml chlorophyll was applied on Quantifoil Au R1.2/1.3, 300 mesh grid and vitrified using Vitrobot (3 sec blot at 4°C and 100% humidity). Data were collected under Titan Krios using EPU 1.9 software on a 300 kV Titan Krios microscope (Thermo Fisher Scientific) equipped with a Gatan Quantum energy filter at a slit width of 20 eV, and a K3 Summit camera (Ametek). Movies were recorded using superresolution mode at a magnification of ×96,000, corresponding to a pixel size of 0.85 Å. A total of 11,027 movies at a total dose of ∼40.5 e^−^/Å^2^ and a dose rate of 20 e^−^/pixel per sec (30 frames), with a defocus range of −1.0 to −2.0 μm. All the final images were binned, which resulted in a pixel size of 0.85 Å for further data processing.

Movies were imported into cryoSPARC^20^ and 1,654,009 particles were boxed using Blob/Template picker. Particles were extracted, and after several rounds of two-dimensional (2D) classification, 268,027 particles were selected for *Ab initio* reconstruction. After two rounds of Hetero-refinement, 88,960 particles were used to construct PSI-SOD-FCPa-FCPb) at 2.66 Å resolution. For further analysis of the peripheral antennae, we use three rounds of Hetero-refinement that yielded a homogeneous class of 56,990 particles that was used to construct PSI-SOD-FCPI at 2.94 Å resolution. Two masks around FCPa-FCPc and FCPd-FCPe were used to improve the local resolution to 3.03 Å and 2.77 Å, respectively. A composite map was generated and used for model building and refinement.

### Model building and refinement

CryFold^21^ was used to build the initial model using available sequence database^51^. Final modelling and real-space refinement were then carried out using *Coot* 0.8.9^22^, particularly the C-terminus of PsaD, PsaL and PsaM, as well as SOD1 and SOD2. All protein residues and pigments were fitted using locally optimized map weights, and restraint files for pigments and ligands were generated using the Grade server (http://grade.globalphasing.org) as previously described^52^. The pigment modeling was consistent with the HPLC analysis. Discrimination of the chromeraxanthin molecule was based on the extended density corresponding to its acetyl group (Extended Data Fig. 4). Compared to typical fucoxanthin found in diatoms^37–40,44^, chromeraxanthin contains a similar acetyl groups, but differs in ring structure. In fucoxanthin, the 3’-hydroxy-5’-keto-cyclohexene (six-membered) ring is intact, whereas in chromeraxanthin it is split into a keto group and a five-membered ring bearing a hydroxyl group. As a result, fucoxanthin adopts a flatter conformation in side view, while chromeraxanthin appears more elliptical. This structural difference aligns better with the observed density map, further supporting the assignment of chromeraxanthin (Extended Data Fig. 9). Discrimination between violaxanthin and Bcr was based on the presence of bulged densities corresponding to the terminal oxygen atom. All Bcr molecules were modeled within PSI core subunits.

The PSI-SOD-FCPI model was refined using Real-Space-Refine from the PHENIX suite^23^. Interactions of validation, model building and refinement were carried out using MolProbity 4.2^53^, *Coot* 0.8.9^22^ and the PHENIX suite^23^. All the buried surfaces were calculated using the online tool PDBePISA v.1.52^54^ (https://www.ebi.ac.uk/pdbe/pisa/). The statistics for cryo-EM data collection, model building and refinement parameters are summarized in the Supplementary Table 1. The correspondence of pigment numbers in FCP described in this study and those of the typical FCPIs is listed in Supplementary Table 5. The figures were prepared using UCSF ChimeraX 1.9^55^.

### Computational analysis of FRET

The FRET rate constant was calculated using the equation: k_FRET = (Cκ^2) / (n^4 R^6), where C is the spectral overlap between the donor fluorescence and the acceptor absorption spectra. For Chl *a* to Chl *a* energy transfer, an empirical value of C = 32.26 was used. is The dipole orientation factor κ^2^ is defined as κ^2 = [(u_D)·(u_A) - 3 ((u_D)·(R_DA)) ((u_A)·(R_DA))] ^2. Where (u_D) and (u_A) are unit vectors from the NB to ND atoms of the donor and acceptor Chl, respectively, representing their transition dipole directions. R_DA is the unit vector from the magnesium atoms of the donor and acceptor Chl. The refractive index ’n’ was set to 1.55, based on prior studies. R is the distance between the magnesium atoms of the donor and acceptor Chls. The FRET lifetime is defined as t_FRET = 1 / k_FRET.

## Data availability

The composite cryo-EM map and atomic coordinates for the PSI-SOD-FCPI supercomplex have been deposited in the Electron Microscopy Data Bank under accession code EMD-##### and the Protein Data Bank under accession code ####. The masked cryo-EM maps have been deposited in the Electron Microscopy Data Bank under accession codes EMD-#####, EMD-#####, and EMD-#####.

Other atomic coordinates that were used in this study: cyanobacterial (PSI PDB ID: 1JB0, https://www.wwpdb.org/pdb?id=pdb_00001jb0), green algae (PDB ID: 7YCA, https://www.wwpdb.org/pdb?id=pdb_00007yca, PDB ID: 7ZQC, https://www.wwpdb.org/pdb?id=pdb_00007zqc), diatom (PDB ID: 8ZEH, https://www.wwpdb.org/pdb?id=pdb_00008zeh), dinoflagellate (PDB ID: 8JJR, https://www.wwpdb.org/pdb?id=pdb_00008jjr), plant (PDB: 8BCV, https://www.wwpdb.org/pdb?id=pdb_00008bcv), PSI-Fd (PDB ID: 6YAC, https://www.rcsb.org/structure/6YAC),

## Acknowledgements

We thank L. Huang, Z. Liu and Y. Zhang from the cryo-EM Facility Westlake University for providing support with data collection, HPC Center of Westlake University for computational resources, R. Sobotka and A. Naschberger for participating in the initial stage of the project, X. Zhang for calculating the energy pathways by FRET, and Z. Li for help with refinement, H. Mu for help with cell culture. This work was supported by the National Key R&D Program of China (2025YFA0921100), the European Research Council (ERC805230), Zhejiang Province Key R&D Program (2023SDXHDX0002), the National Key Laboratory of Gene Expression, Zhejiang Key Laboratory of Low-Carbon Intelligent Synthetic Biology (2024ZY01025), Westlake Research Center for Industries of the Future (WU2023C002), and the Westlake Center for Genome Editing.

## Author contributions

Y.F. performed the experiments, model building, analysis, and wrote the paper. K.H. performed data collection, structure determination, and contributed to the model building, analysis, and writing the paper. X.L. supervised the project. A.A. analyzed the structure, wrote the paper, and supervised the project. All authors contributed to the analysis and the final version of the paper.

## Competing interests

The authors declare no competing interests.

---

**Extended Data Fig. 1.**
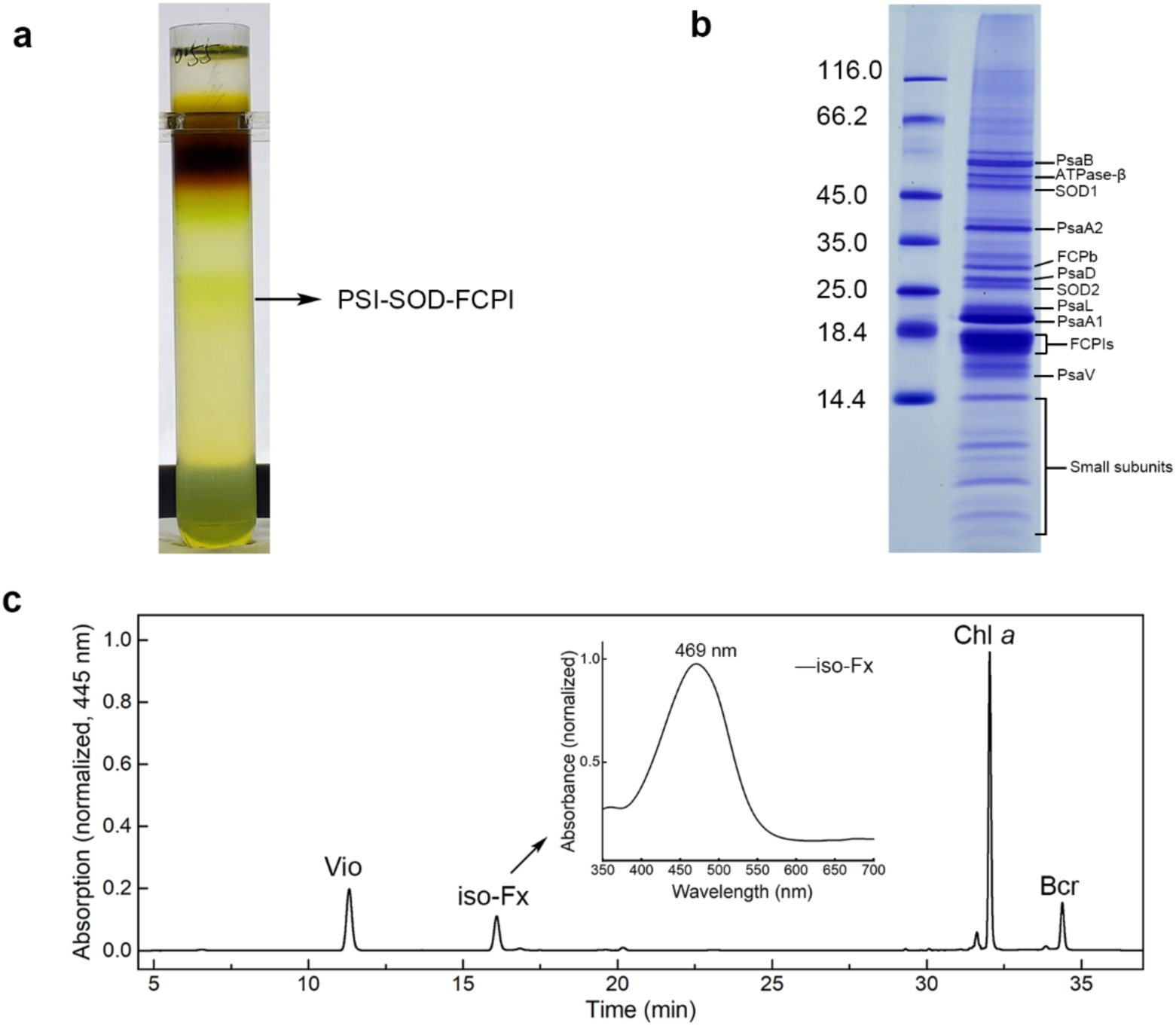
Sample preparation and characterization. **a,** Sucrose density gradient of purified PSI-SOD-FCPI. The PSI fraction is indicated. **b,** SDS-PAGE analysis of the purified PSI-SOD-FCPI (5 μg Chl *a*). The presence of small protein subunits was confirmed by mass spectrometry analysis. **c**, Analysis of the pigments of PSI-SOD-FCPI by HPLC monitored at 445 nm. Four major pigment peaks were identified, including Chlorophyll *a* (Chl *a*), iso-fucoxanthin (iFx, chromeraxanthin), violaxanthin (Vio), and β-carotene (Bcr). The absorption spectrum of the chromeraxanthin pigment was measured at room temperature.

**Extended Data Fig. 2.**
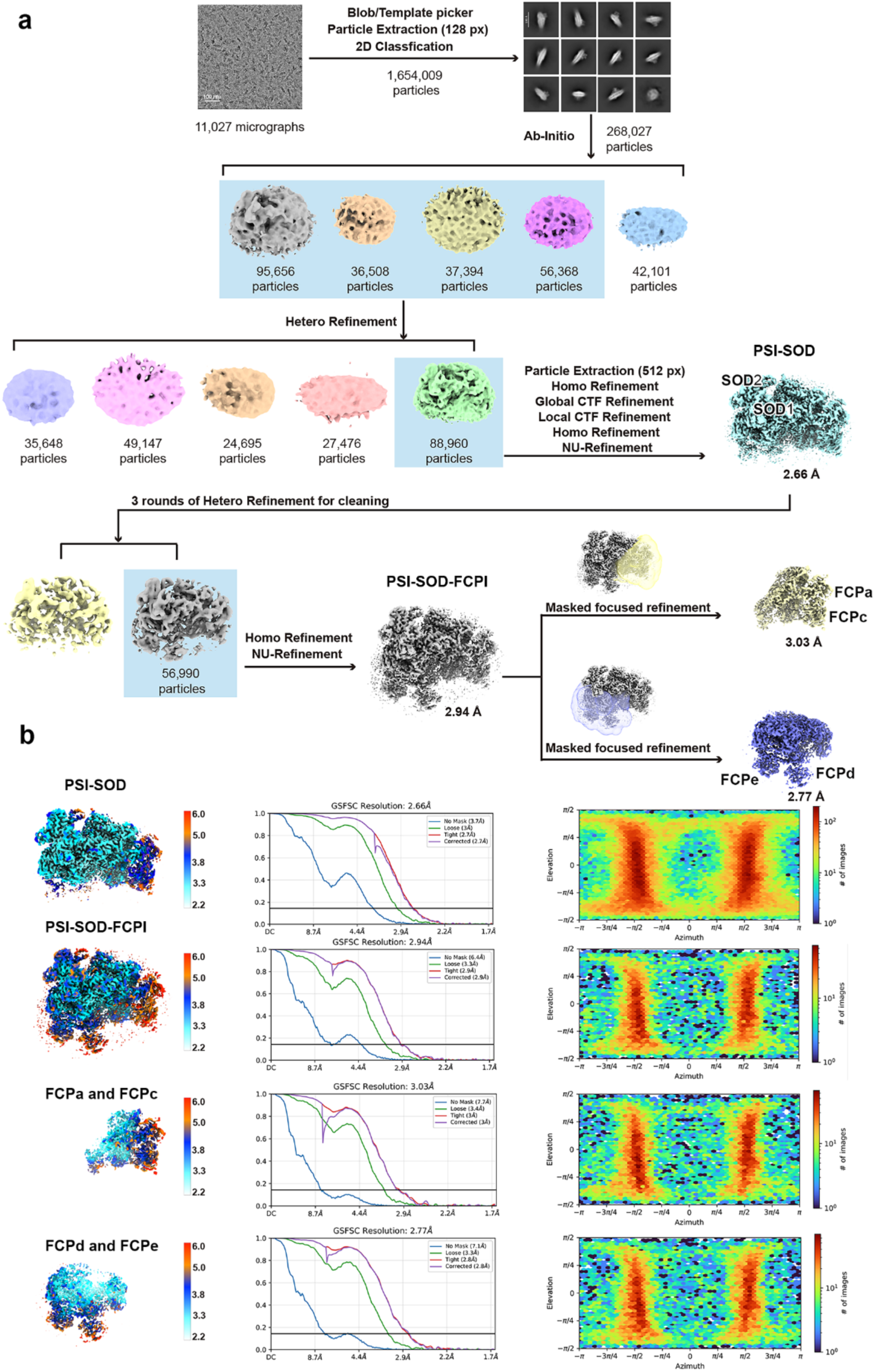
Cryo-EM processing workflow and local resolution. **a,** Flowchart of the data processing for the PSI-SOD-FCPI supercomplex and antenna proteins. **b,** The maps of the PSI-SOD, PSI-SOD-FCPI and masked FCP light-harvesting proteins are shown colored by local resolution (left) with their corresponding FSC curves and angular distribution.

**Extended Data Fig. 3.**
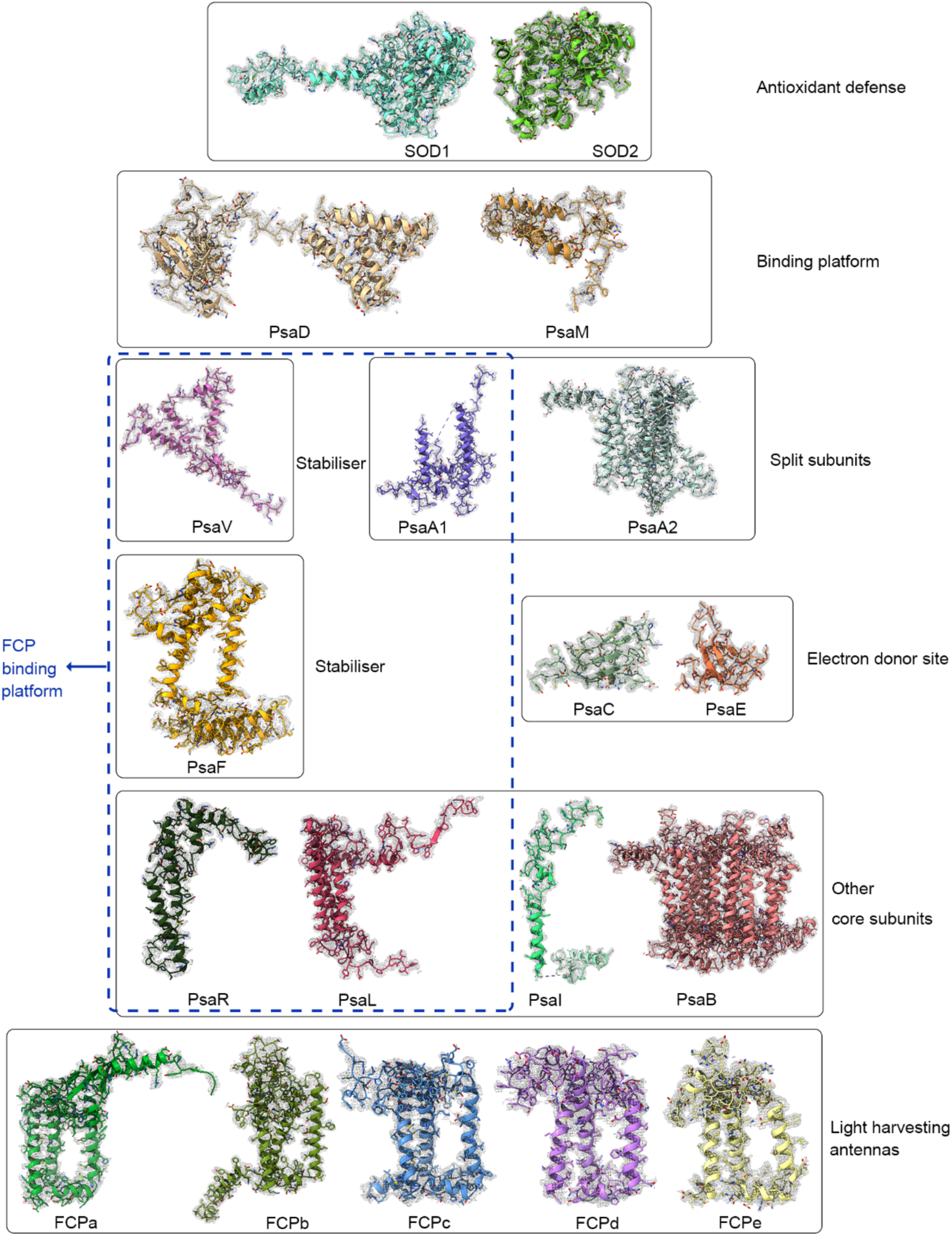
Individual subunits with their cryo-EM density maps. Protein subunits are shown in stick and cartoon representation with the N and O atoms in blue and red, respectively. The density maps are shown with a threshold of 0.14 contour level (step 1) in mesh by Volume Viewer in ChimeraX 1.9^55^.

**Extended Data Fig. 4.**
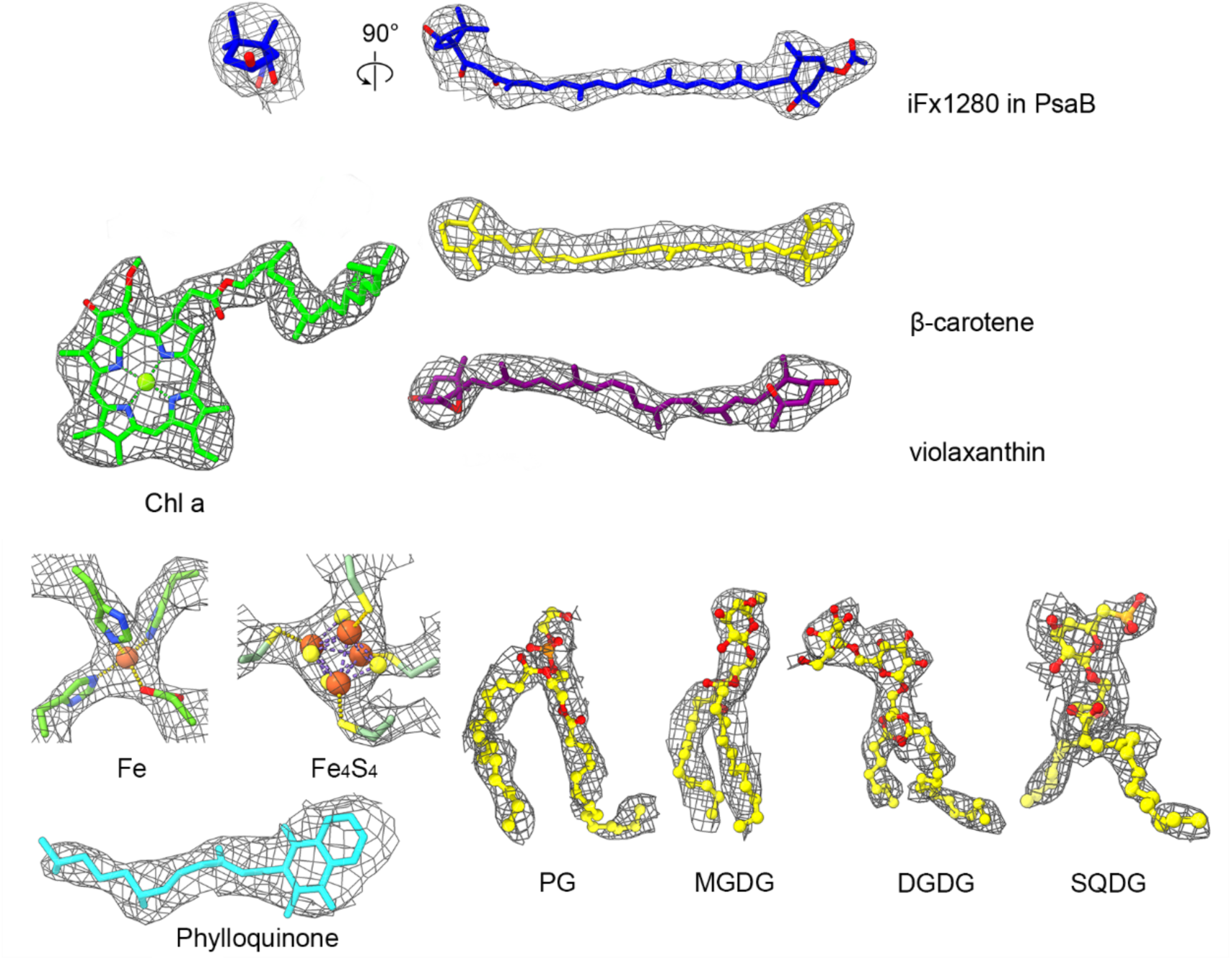
Examples of endogenous cofactors and lipids identified in the map.

**Extended Data Fig. 5.**
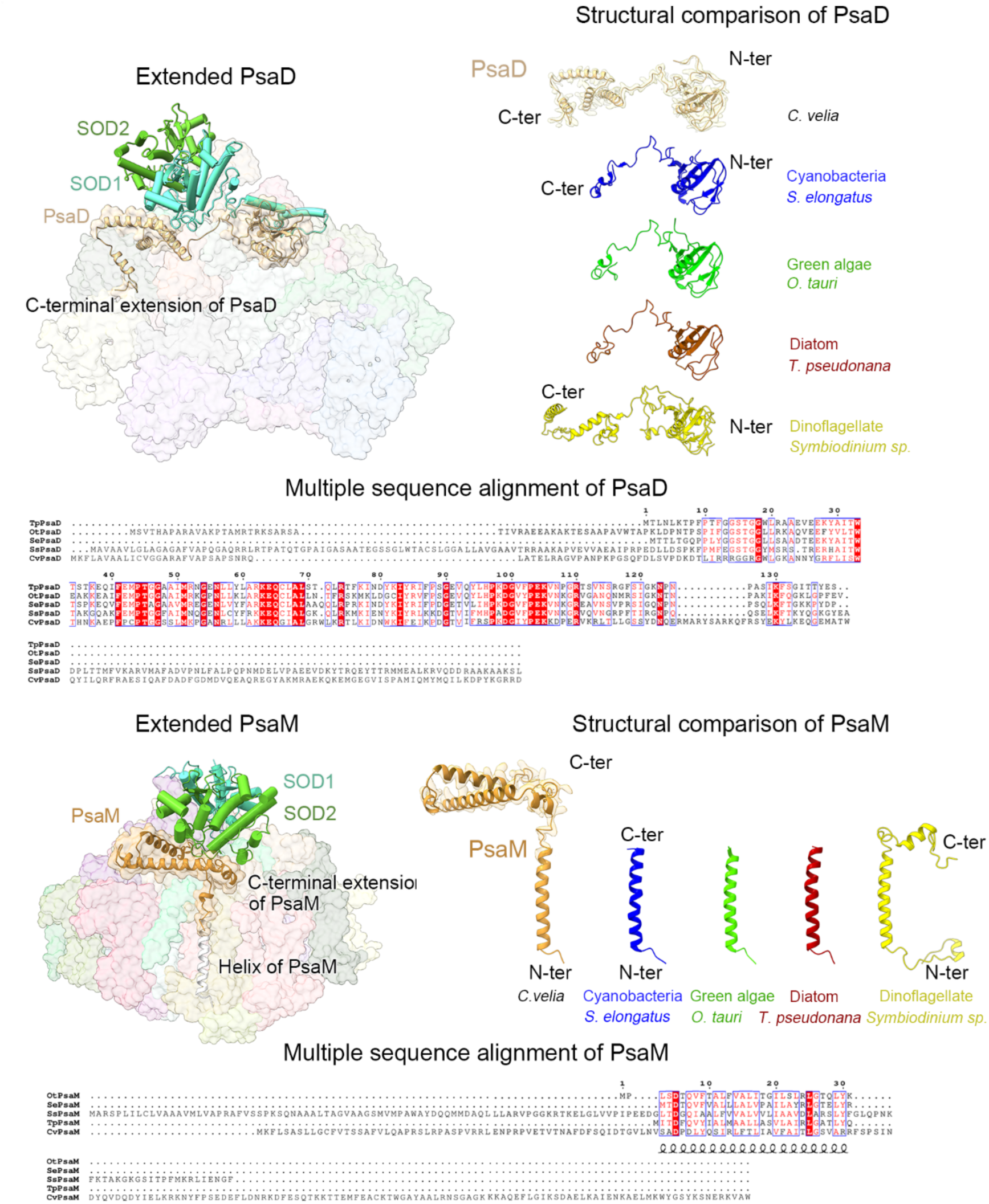
Comparison of the PsaD and PsaM subunits in PSI from different organisms. PsaD and PsaM of *C. velia* are more extended at the C-terminus in the stroma in comparison to cyanobacteria (PDB ID: 1JB0, https://www.wwpdb.org/pdb?id=pdb_00001jb0), green algae (PDB ID: 7YCA, https://www.wwpdb.org/pdb?id=pdb_00007yca), diatoms (PDB ID: 8ZEH, https://www.wwpdb.org/pdb?id=pdb_00008zeh) and dinoflagellates (PDB ID: 8JJR, https://www.wwpdb.org/pdb?id=pdb_00008jjr).

**Extended Data Fig. 6.**
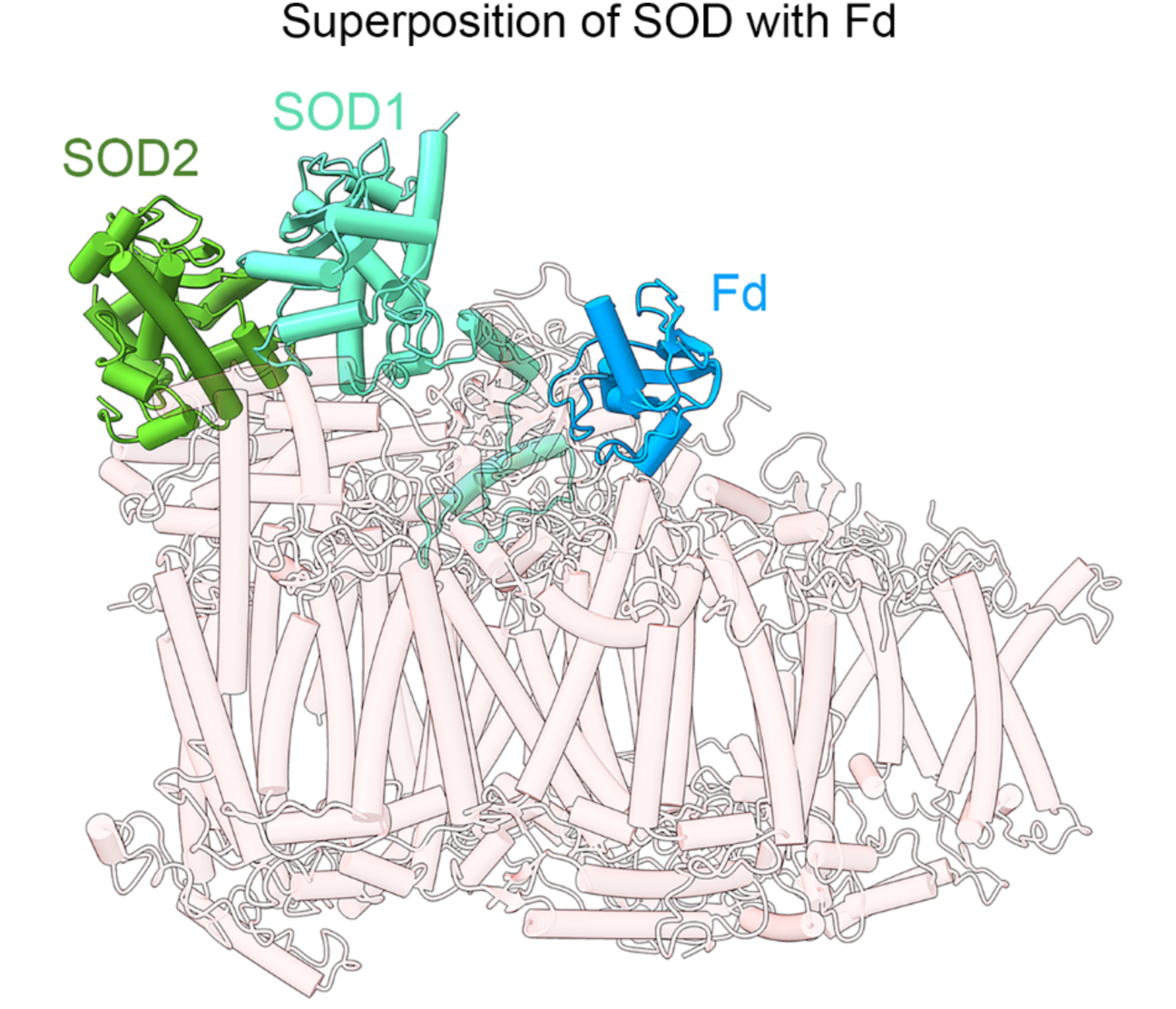
Structural superposition of PSI-SOD-FCPI with plant PSI-bound ferredoxin. Superposition of PSI-SOD model with ferredoxin from pea PSI-Fd model^25^ (PDB ID: 6YAC, https://www.rcsb.org/structure/6YAC) displays no steric clashes between SOD and ferredoxin.

**Extended Data Fig. 7.**
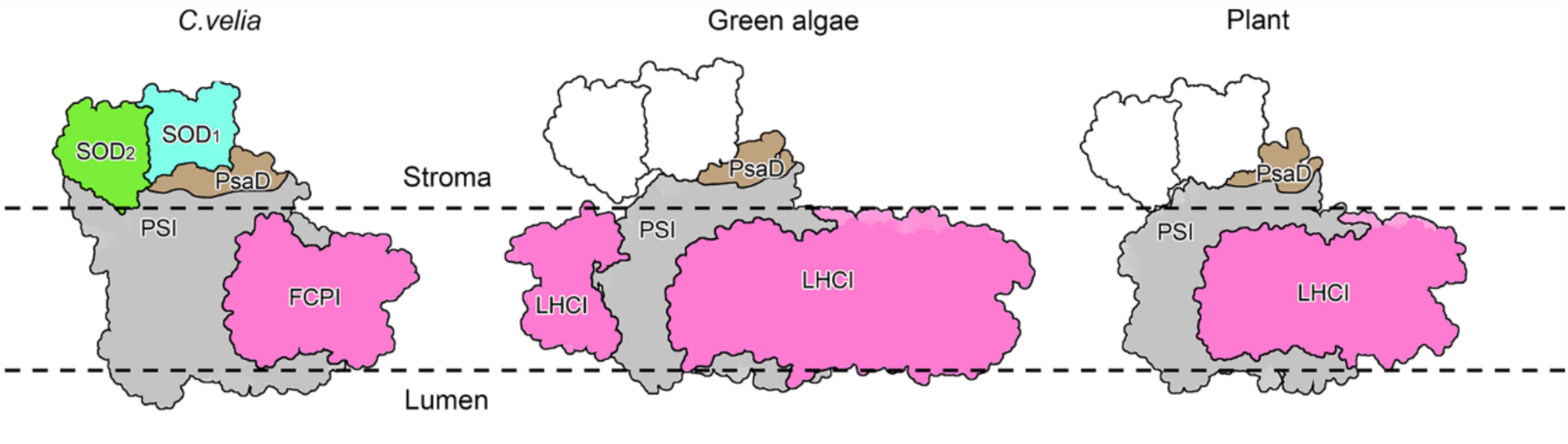
Comparison of SOD’s relative position at the green algae and plant PSI interface. Left, surface representation of the *C. velia* PSI-SOD-FCPI. Middle and right, schematic representation of green algae ^29^ (PDB ID: 7ZQC, https://www.wwpdb.org/pdb?id=pdb_00007zqc) and plant ^27^ (PDB: 8BCV, https://www.wwpdb.org/pdb?id=pdb_00008bcv) PSI-LHCI, respectively, with the relative position of SOD1, SOD2 outlined.

**Extended Data Fig. 8.**
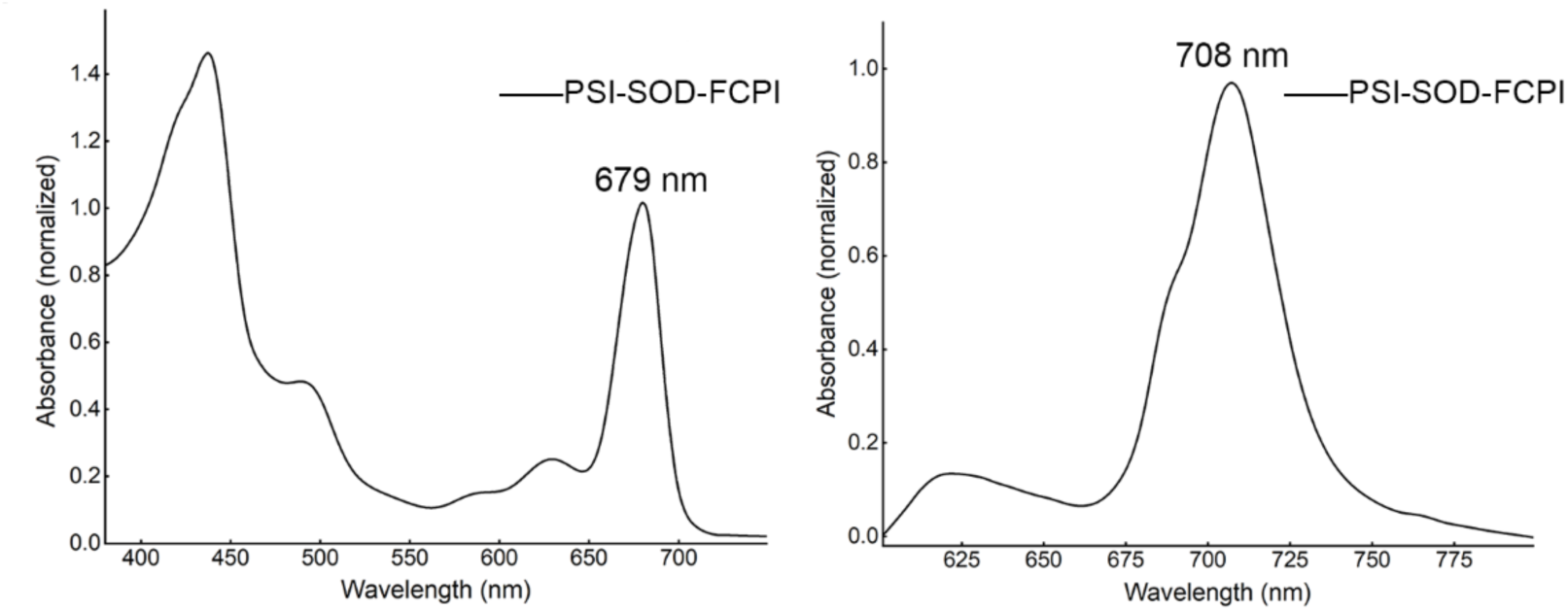
Room temperature absorption spectrum and 77 K fluorescence emission spectra of PSI-SOD-FCPI.

**Extended Data Fig. 9.**
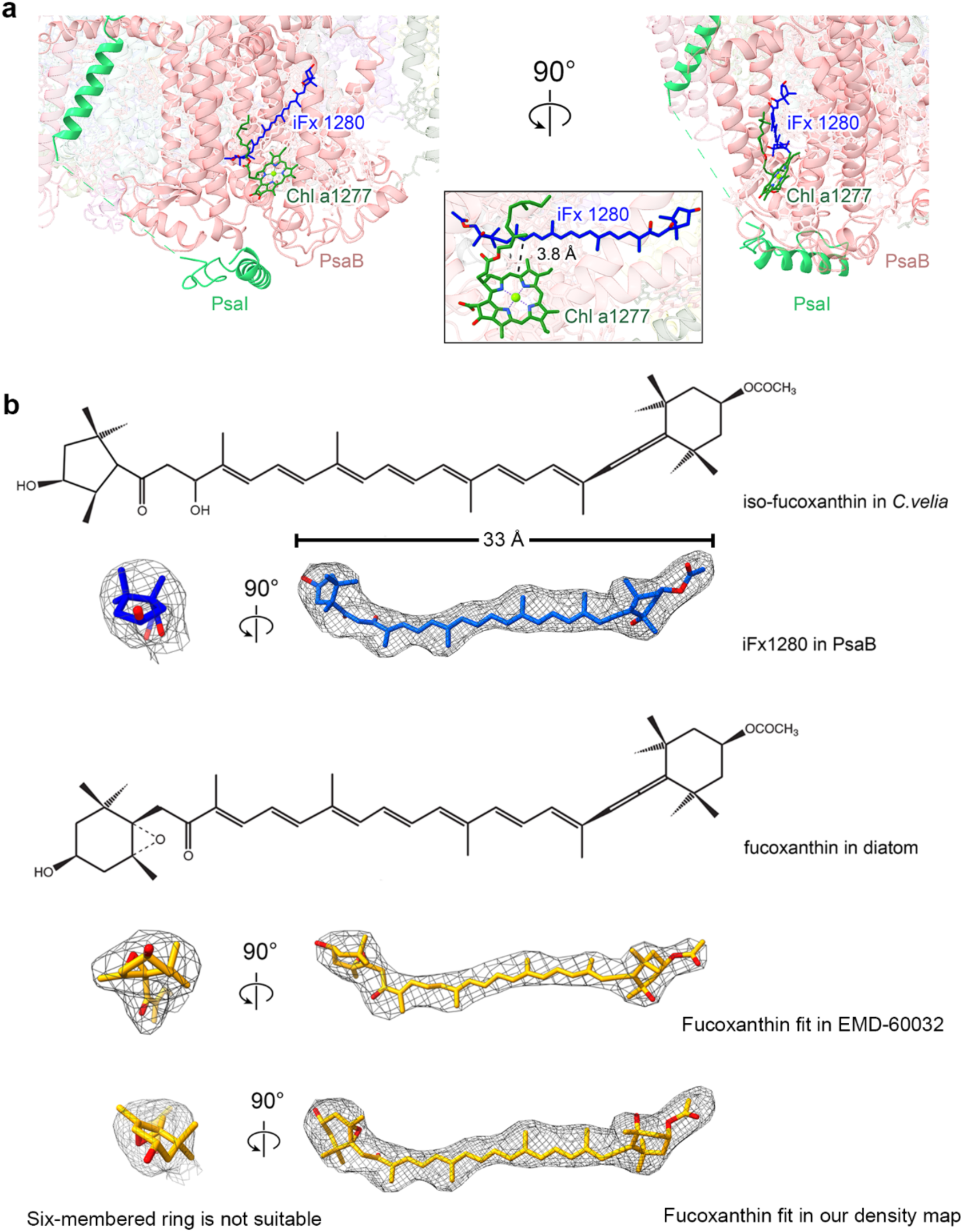
Chromeraxanthin in *C. velia* PSI. **a,** Chromeraxanthin is coordinated by PsaB subunit close to PSI lumenal extension and associated with Chl *a*1277. **b,** The 33-Å density is modeled with chromeraxanthin (blue), which differs from canonical fucoxanthin (yellow) by featuring a cyclopentane ring instead of a cyclohexene ring, as illustrated in the schematic. For comparison, the cyclohexene ring density from EMD-60032 (https://www.emdataresource.org/EMD-60032)^38^ is shown, exhibiting a flat density that does not accommodate the cyclopentane isoform observed here (bottom panel).

**Extended Data Fig. 10.**
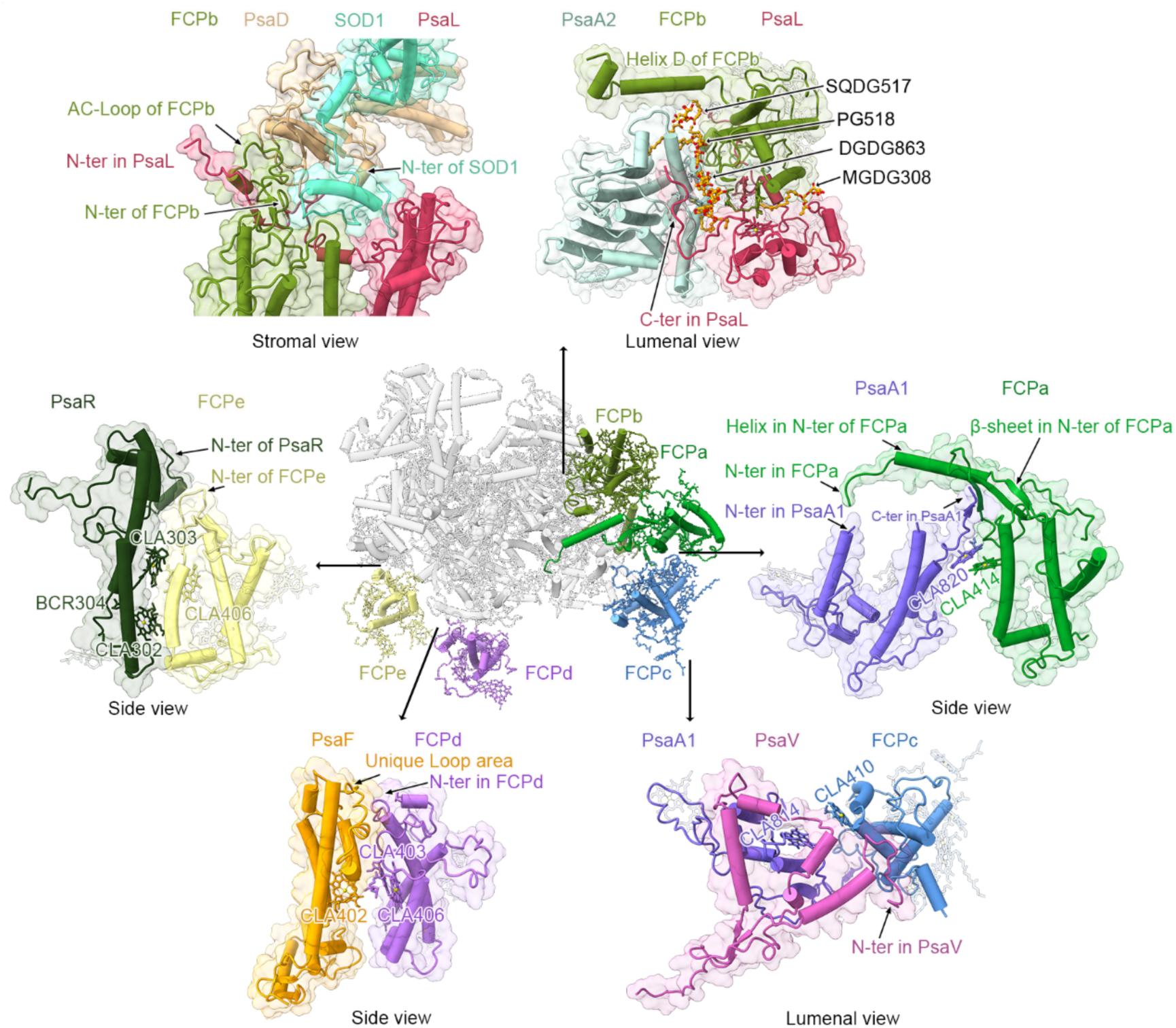
Structural adaptation of FCP proteins to the PSI core surface. Each FCP protein shares a conserved central domain comprising three transmembrane helices but exhibits unique peripheral adaptations to interact with distinct PSI core subunits. FCPa features an N-terminal extension that engages the C-terminus of PsaA1. FCPb has an N-terminal loop that interacts with PsaD and PsaL, while its C-terminal helix extends toward PsaA2. FCPc has co-evolved to associate with the N-terminus of PsaV on the lumenal side. FCPd binds near the loop region of PsaF, and FCPe interacts with PsaR. These structural variations result in distinct orientations of individual FCP proteins relative to the core and establish different sets of functionally coupled chlorophylls (labeled) with PSI subunits.

**Extended Data Fig. 11.**
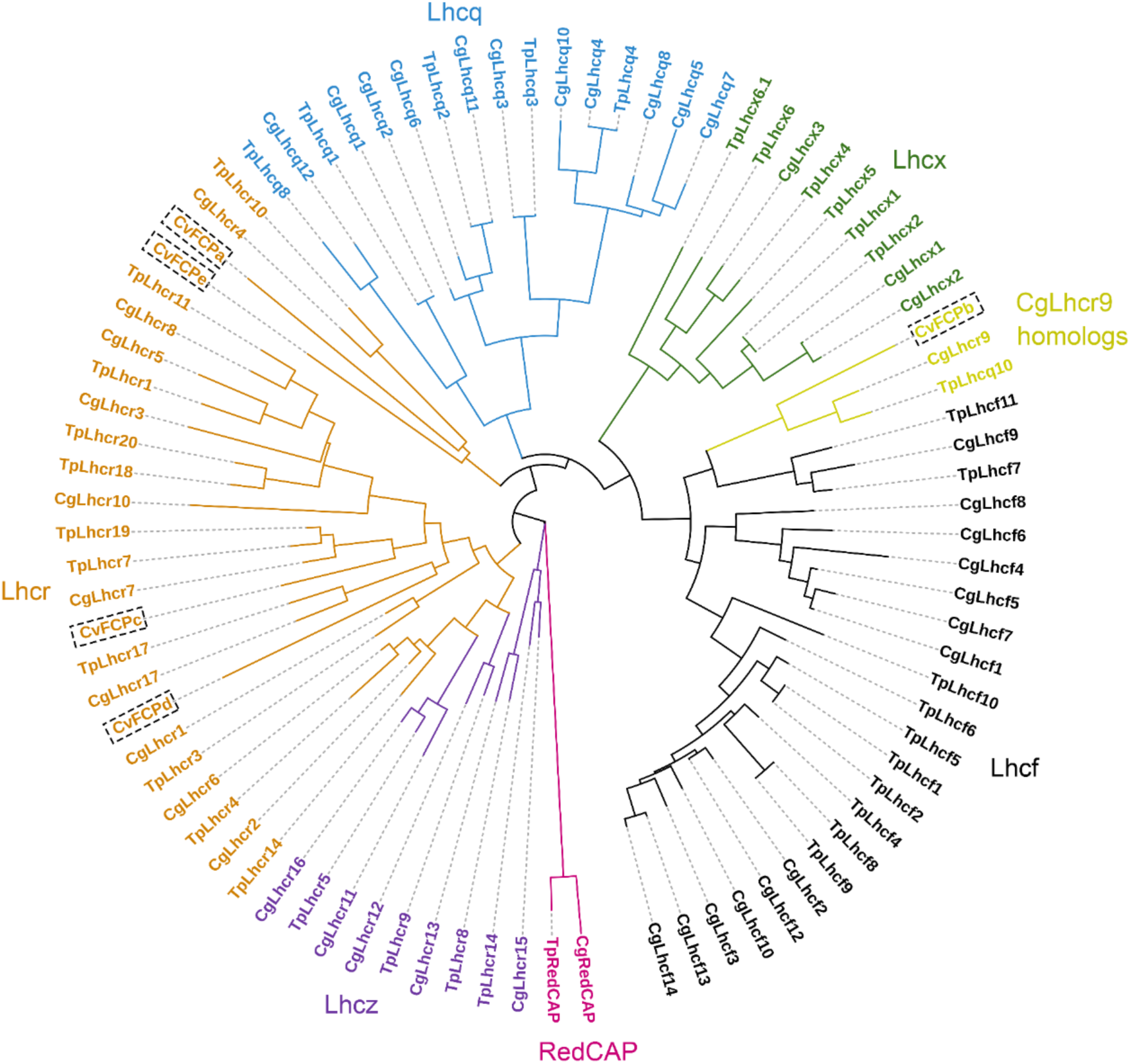
Phylogenetic analysis of FCP proteins. Phylogenetic tree of FCP proteins characterized in this study alongside representative diatom FCPs, including Lhcf, Lhcr, Lhcx, Lhcq, Lhcz and RedCAP family members from *T. pseudonana* and *C. gracilis*. The FCPs identified in this study are highlighted. RedCAP family pink, Lhcf black, Lhcq blue, Lhcr orange, Cg-Lhcr9 yellow, Lhcx green.

**Extended Data Fig. 12.**
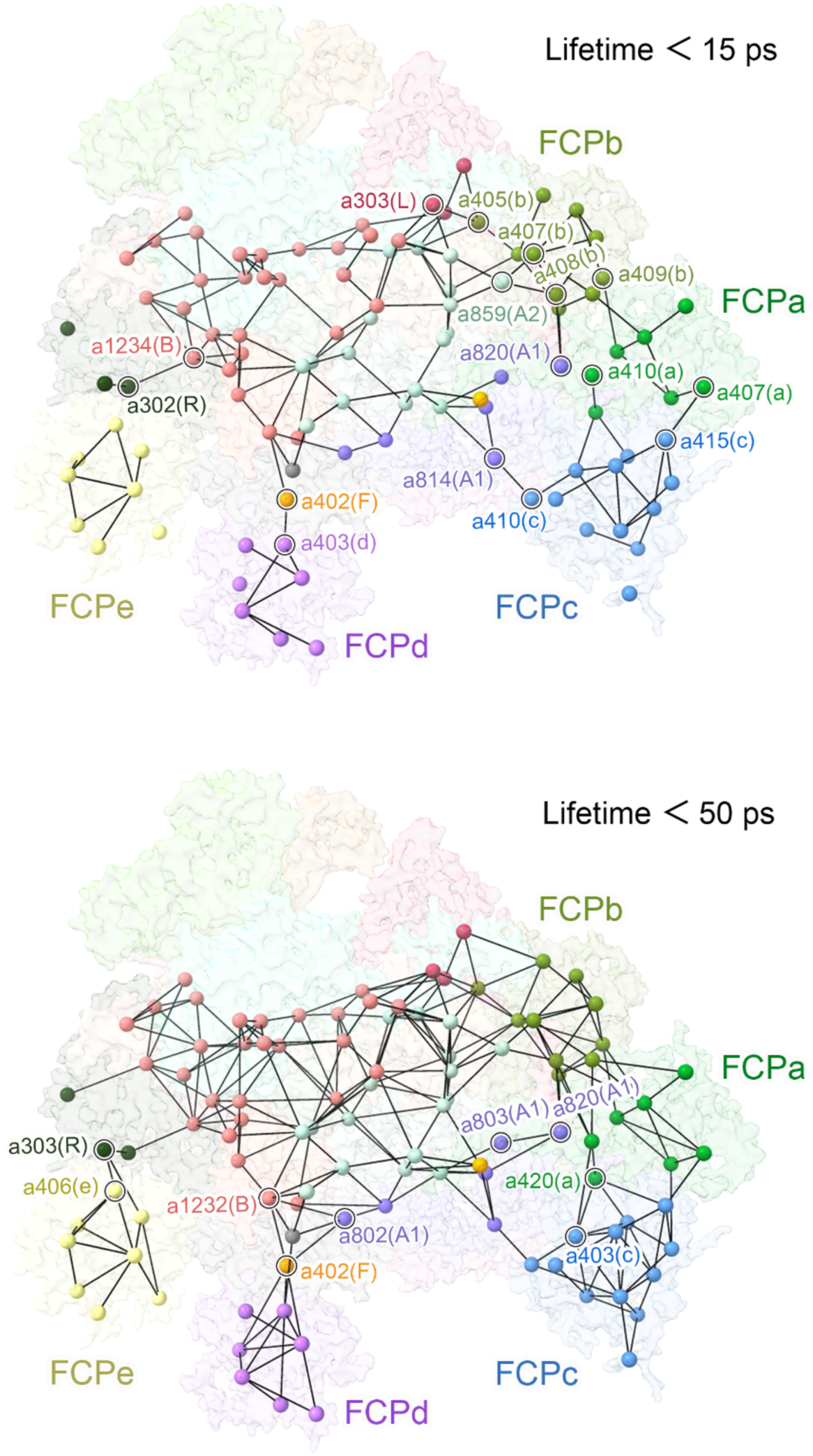
Structure-based analysis of FRET networks within the PSI-SOD-FCPI supercomplex. **a,** Organization of Chls *a* within the PSI-SOD-FCPI supercomplex highlighting FRET pathways. Networks with energy transfer lifetimes under 15 ps and 50 ps are shown. The Chls *a* are represented as spheres and are color-coded by subunit.

**Supplementary Information Table 1.** Cryo-EM data collection, refinement, and validation statistics of PSI-SOD-FCPI.

| PSI-SOD-FCPI (PDB: #####; EMD-#####) |  |
| --- | --- |
| **Data collection and processing** |  |
| Magnification | 96,000 |
| Voltage (kV) | 300 |
| Electron exposure (e<sup>-</sup>/Å<sup>2</sup>) | 50 |
| Defocus range(μm) | -0.6~-2.4 |
| Initial images (no.) | 11,027 |
| Symmetry imposed | C<sub>1</sub> |
| Final particle images(no.) | 56,990 |
| Map resolution(Å) | 2.94 |
| **Refinement** |  |
| Initial model used | CryFold |
| Map sharpening b factor (Å<sup>2</sup>) | -21.23 |
| Model composition |  |
| Non-hydrogen atoms | 39892 |
| Protein residues | 4296 |
| Ligands | 135 |
| B factors (Å<sup>2</sup>) (min/max/mean) |  |
| Protein | 25.56/332.06/107.88 |
| Ligand | 48.43/217.24/110.19 |
| R.m.s.deviation |  |
| Bond lengths (Å) | 0.003 |
| Bond angles (Å) | 0.930 |
| Validation |  |
| MolProbity score | 2.29 |
| Clashscore | 6.08 |
| Poor rotamers (%) | 3.45 |
| Ramachandran plot |  |
| Favored (%) | 96.23 |
| Allowed (%) | 3.70 |
| Disallowed (%) | 0.07 |

**Supplementary Information Table 2.** Lengths of peptides and their interacting cofactors in the structure of PSI-SOD-FCPI.

| Protein | Chain<br>in PDB | Source<br>(UniProt) | Residues | Pigments | Lipids and ligands |
| --- | --- | --- | --- | --- | --- |
| PsaA1 | a | D9IXH7 | 35-122,<br>141-240 | 6 Chls *a* |  |
| PsaA2 | v | D9IXI2 | 1-395 | 18 Chls *a*, 3<br>Bcrs | 1 phylloquinone,<br>1 Fe<sub>4</sub>S<sub>4</sub>,<br>1 DGDG, 1 MGDG |
| PsaB | b | D9IXI4 | 324-482,<br>560-1139 | 28 Chls *a*, 4<br>Bcrs,<br>1 iFx | 1 phylloquinone<br>2 PG |
| PsaC | c | D9IXG0 | 1-80 |  | 2 Fe<sub>4</sub>S<sub>4</sub> |
| PsaD | d | A0A0G4H942 | 36-268 |  |  |
| PsaE | e | A0A0G4H6U3 | 45-110 |  |  |
| PsaF | f | A0A0G4GX86 | 103-326 | 2 Chls *a* |  |
| PsaI | i | A0A0G4HFQ7 | 86-122,<br>151-211 |  |  |
| PsaL | l | A0A0G4GX71 | 52-277 | 3 Chls *a*, 3 Bcrs | 1 MGDG |
| PsaR | r | A0A0G4FDJ8 | 42-200 | 3 Chls *a*, 1 Bcr |  |
| SOD1 | o | A0A0G4HJZ8 | 59-346 |  |  |
| SOD2 | q | A0A0G4FB02 | 47-266 |  | 1 Fe |
| PsaM | m | A0A6U4Q677 | 86-195 |  |  |
| PsaV | x | A0A0G4HRT4 | 279-423 |  |  |
| FCPa | U | A0A0G4I0G6 | 24-65,<br>92-270 | 7 Chls *a* |  |
| FCPb | G | A0A0G4ID45 | 46-293 | 10 Chls *a*, 4<br>XATs | 1 SQDG |
| FCPc | T | A0A0G4FL36 | 42-236 | 12 Chls *a*, 1<br>XAT |  |
| FCPd | N | A0A0G4HMN0 | 47-212 | 7 Chls *a*, 1 XAT |  |
| FCPe | K | A0A0G4HNB1 | 56-216 | 8 Chls *a*, 1 XAT |  |
| Total of PSI-SOD-FCPI supercomplex |  |  |  | 104 Chls *a*,<br>11 Bcrs,<br>1 iFx, 7 XATs | 2 phylloquinones,<br>3 Fe<sub>4</sub>S<sub>4</sub>, 1 Fe<br>1 DGDG, 2<br>MGDG,<br>2 PG, 1 SQDG |

**Chl *a***: chlorophyll *a*; **Bcr**: β-carotene; **iFx**: iso-fucoanthin (chromeraxanthin); **XAT**: violaxanthin;
**DGDG**: digalactosyl diacyl glycerol; **MGDG**: 1,2-distearoyl-monogalactosyl-diglyceride; **PG**: 1,2-Dipalmitoyl-phosphatidyl-glycerol; **SQDG**: 1,2-di-O-acyl-3-O-[6-deoxy-6-sulfo-α-D-glucopyranosyl]-Sn-glycerol.

**Supplementary Information Table 3.**
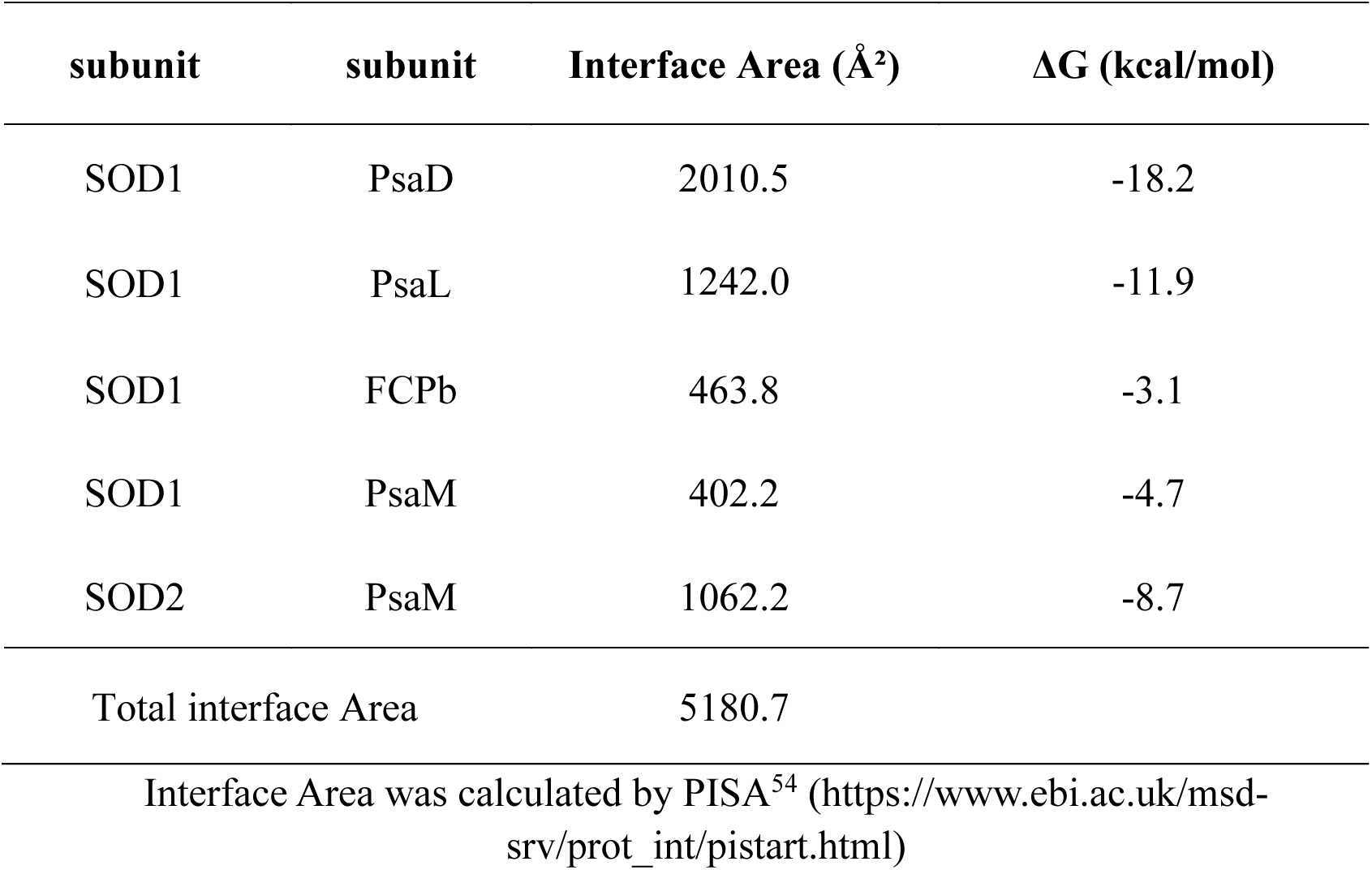
Interface Area (Å²) between SOD1/2 with PSI-core subunits and FCPb in PSI-SOD-FCPI.

**Supplementary Information Table 4.** Calculated FRET rates of the main pathway within the PSI-SOD-FCPI supercomplex.

| Energy pathway in pigment pairs | FRET rate, $k_{FRET}$ (ps) | Lifetime, $\tau$ (ps) | Distance, $R_{DA}$ (Å) | Dipole orientation factor, $\kappa^2$ |
| --- | --- | --- | --- | --- |
| Chl *a*407<sub>FCPb</sub>→Chl *a*859<sub>PsaA2</sub> | 1.052 | 0.951 | 11.102 | 0.352 |
| Chl *a*408<sub>FCPb</sub>→Chl *a*859<sub>PsaA2</sub> | 0.231 | 4.337 | 16.245 | 0.758 |
| Chl *a*405<sub>FCPb</sub>→Chl *a*303<sub>PsaL</sub> | 0.232 | 4.316 | 19.572 | 2.33 |
| Chl *a*408<sub>FCPb</sub>→Chl *a*820<sub>PsaA1</sub> | 0.18 | 5.551 | 18.538 | 1.308 |
| Chl *a*409<sub>FCPb</sub>→Chl *a*410<sub>FCPa</sub> | 0.28 | 3.571 | 19.974 | 3.182 |
| Chl *a*407<sub>FCPa</sub>→Chl *a*415<sub>FCPc</sub> | 0.286 | 3.497 | 16.93 | 1.205 |
| Chl *a*410<sub>FCPc</sub>→Chl *a*814<sub>PsaA1</sub> | 0.609 | 1.642 | 14.687 | 1.094 |
| Chl *a*403<sub>FCPd</sub>→Chl *a*402<sub>PsaF</sub> | 0.408 | 2.451 | 12.221 | 0.243 |
| Chl *a*820<sub>PsaA1</sub>→Chl *a*803<sub>PsaA1</sub> | 0.047 | 21.137 | 19.293 | 0.436 |
| Chl *a*420<sub>FCPa</sub>→Chl *a*403<sub>FCPc</sub> | 0.12 | 18.349 | 19.375 | 1.134 |
| Chl *a*402<sub>PsaF</sub>→Chl *a*802<sub>PsaA1</sub> | 0.06 | 16.738 | 21.454 | 1.042 |
| Chl *a*402<sub>PsaF</sub>→Chl *a*1232<sub>PsaB</sub> | 0.082 | 16.152 | 18.18 | 0.532 |
| Chl *a*406<sub>FCPe</sub>→Chl *a*303<sub>PsaR</sub> | 0.047 | 21.421 | 11.511 | 0.019 |
| Chl *a*303<sub>PsaR</sub>→Chl *a*302<sub>PsaR</sub> | 0.123 | 8.148 | 18.259 | 0.814 |
| Chl *a*302<sub>PsaR</sub>→Chl *a*1234<sub>PsaB</sub> | 0.087 | 11.446 | 19.202 | 0.784 |

**Supplementary Information Table 5.**
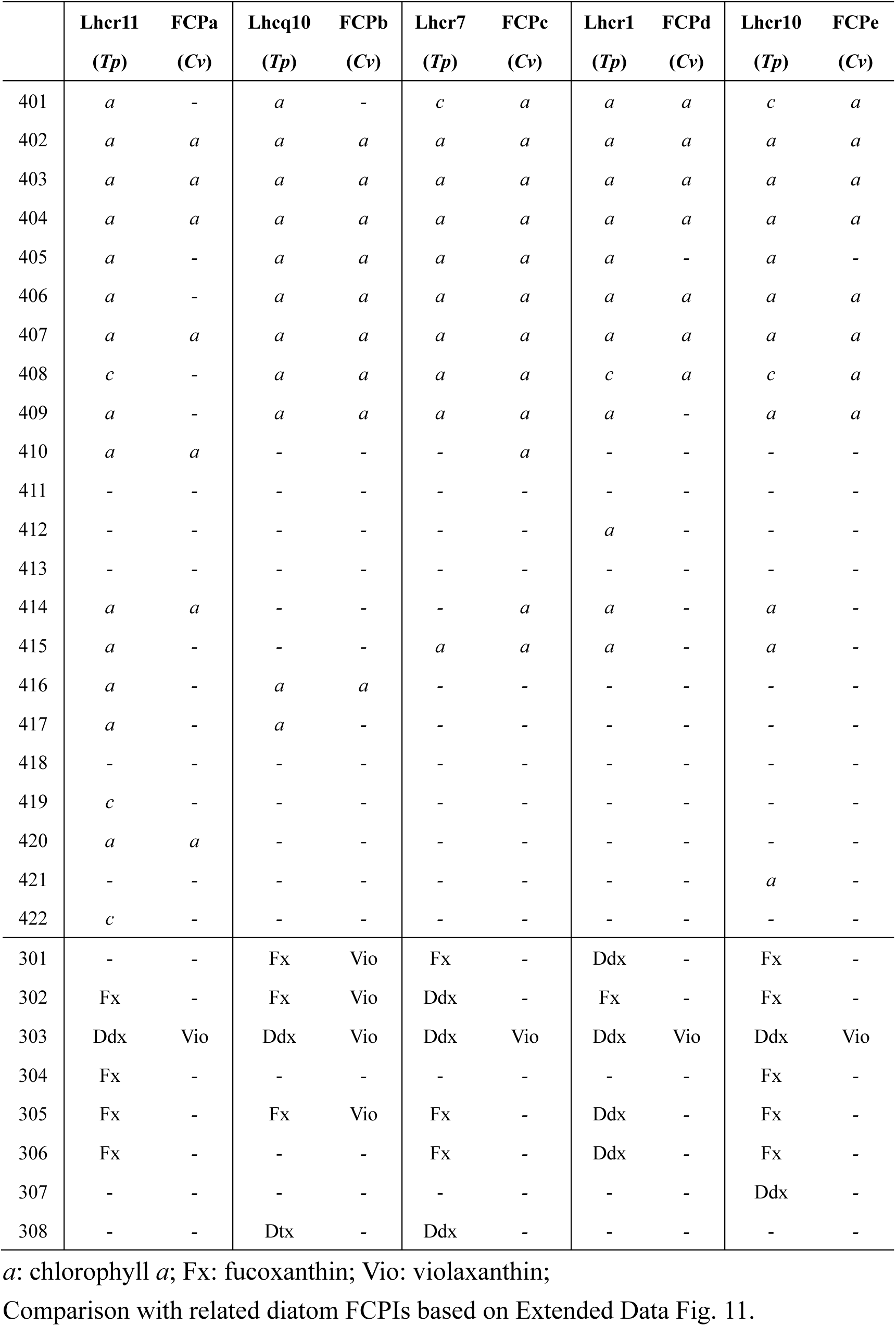
Pigment-binding sites of the 5 FCPIs in PSI-SOD-FCPI supercomplex.

## Notes

### Competing Interest Statement

The authors have declared no competing interest.

